# A novel and effective Cre/*lox*-based genetic tool for repeated, targeted and markerless gene integration

**DOI:** 10.1101/2020.12.30.424803

**Authors:** Qinghua Zhou, Liangcheng Jiao, Wenjuan Li, Zhiming Hu, Yunchong Li, Houjin Zhang, Li Xu, Yunjun Yan

## Abstract

The unconventional yeast *Yarrowia lipolytica* is extensively applied in bioproduction fields owing to its excellent metabolite and protein production ability. Nonetheless, utilization of this promising host is still restricted by limited availability of precise and effective gene integration tools. In this study, a novel and efficient genetic tool was developed for targeted, repeated, and markerless gene integration based on Cre/*lox* site-specific recombination system. The developed tool required only a single selection marker and could completely excise all of the unnecessary sequences. A total of three plasmids were created and seven rounds of marker-free gene integration were examined in *Y. lipolytica*. All the integration efficiencies remained above 90%, and analysis of protein production and growth characteristics of the engineered strains confirmed that genome modification via the novel genetic tool was feasible. Further work also confirmed the genetic tool was effective for integration of other genes, loci, and strains. Thus, this study significantly promotes the application of Cre/*lox* system and presents a powerful tool for genome engineering in *Y. lipolytica*.

**Graphical abstract:** 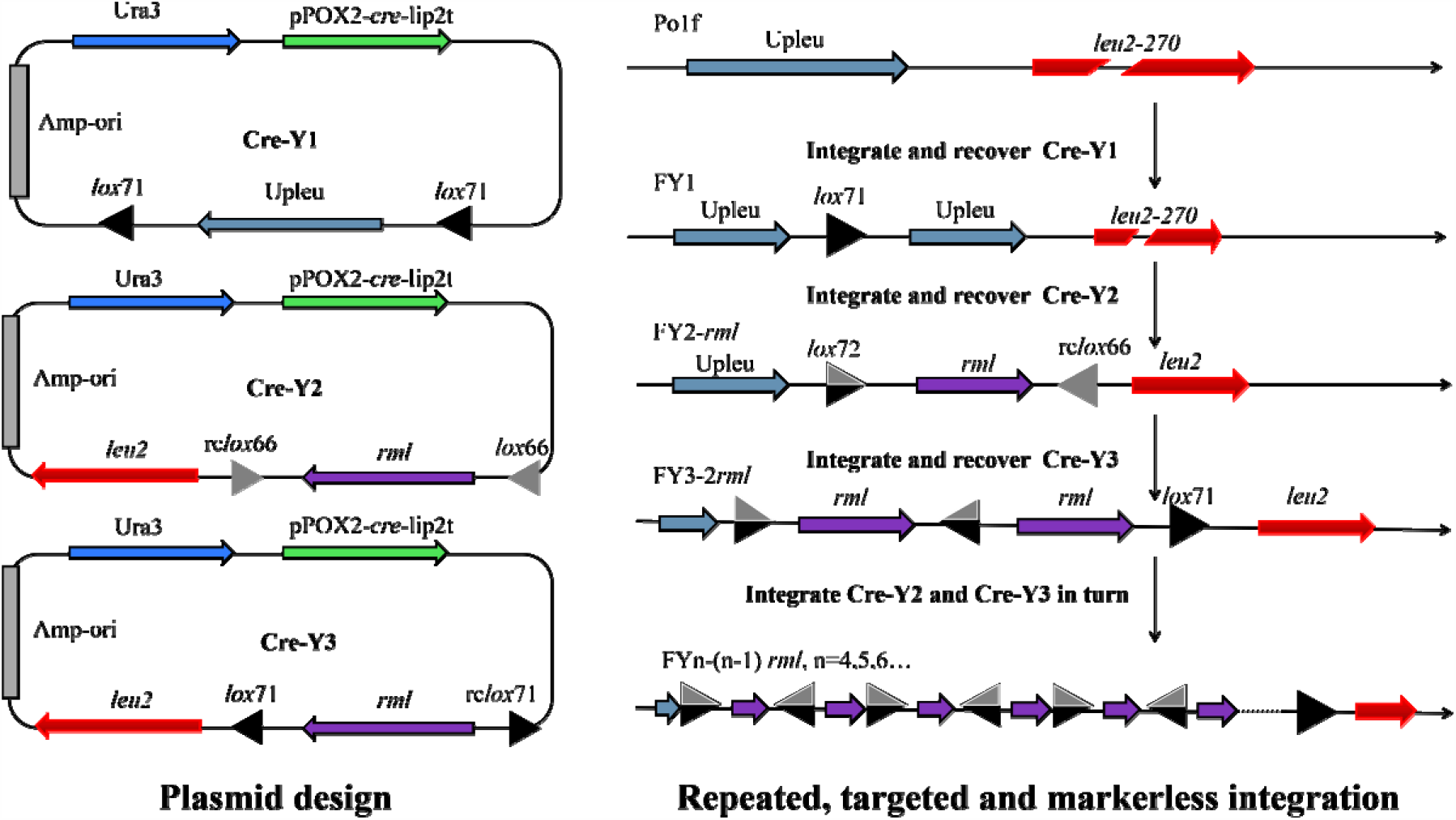

The novel genetic tool comprised three plasmids, namely, Cre-Y1, Cre-Y2, and Cre-Y3. Cre-Y1 was introduced into the Upleu locus of Po1f, and then recombination between two *lox*71 sites could remove unnecessary plasmid elements and produce an initial strain harboring a *lox*71 site. Subsequently, Cre-Y2 was integrated onto the *leu2* locus, forming a transitional strain with three *lox* sites (*lox*71, *lox*66, and rc*lox*66). Recombination reactions among these three *lox* sites could excise unnecessary DNA fragments and retain a *lox*72 site, an rc*lox*66 site, and a target gene expression cassette in the genome. Similarly, integration of Cre-Y3 produced strains harboring two target gene expression cassettes. Consequently, repeated, targeted, and markerless gene integration could be efficiently realized following iterative integration of Cre-Y2 and Cre-Y3.

**Highlights:**

- A powerful genetic tool was developed for targeted and markerless gene integration
- The genetic tool was effective for various genes, loci, and strains
- Only a single selection marker was adequate for repeated gene integration
- All of the useless and redundant sequences would be excised
- All integration efficiencies could remain above 90%

## 1. Introduction

With the development of synthetic biology, various molecular genetic tools are being increasingly designed for precise genome modification in yeasts. As a promising host, the generally regarded as safe (GRAS) yeast, *Yarrowia lipolytica*, has been employed in a wide range of fields such as protein production, metabolic engineering, degradation of *n*-alkanes, and dimorphism (Abdel-Mawgoud et al., 2018; Blazeck et al., 2014; Carly et al., 2017; Czajka et al., 2018; Madzak, 2015; Madzak, 2018). Nevertheless, only few effective genetic tools have been developed for repeatedly targeted gene integration in this host, which still restricts further exploitation and application of *Y. lipolytica*.

In traditional methods, heterologous and endogenous genes are transformed into *Y. lipolytica* genome via insertion of the whole plasmid at target loci or integration of tandem fragment of target gene and selection marker at zeta loci (Barth and Gaillardin, 1996; Larroude et al., 2018; Pignede et al., 2000). However, introduction of non-target DNA fragments, especially antibiotics resistance genes, may harm the GRAS status of *Y. lipolytica* and hinder its application in food and medicine industries. Besides, the bioproduction of *Y. lipolytica* usually requires multi-round integrations of target gene or co-expression of various genes (Carly et al., 2017; Larroude et al., 2018; Ledesma-Amaro and Nicaud, 2016; Markham et al., 2018; Yang et al., 2019), which can be hardly realized via these conventional methods owing to the limited availability of selection markers. Consequently, there is an urgent need to develop efficient genetic tool(s) for targeted, repeatable, and markerless gene integration in *Y. lipolytica*.

To date, several site-specific nuclease technologies have been established for genome modification in various species (Cong et al., 2013; Gaj et al., 2013; Hsu et al., 2014; Wu et al., 2014). Among them, CRISPR-Cas9 system can accomplish relatively high-efficiency gene disruption and marker-free gene integration in *Y. lipolytica* (Bae et al., 2020; Holkenbrink et al., 2018; Schwartz et al., 2017a). A previous study examined 17 loci for targeted and markerless gene integration in *Y. lipolytica*, among which only five loci exhibited gene integration, with highest integration efficiency of 68.9%±25.5% (Schwartz et al., 2017b). Another study accomplished gene integration into *PEX*10 locus via homologous recombination (HR) and homology-mediated end-joining, with integration efficiencies of 6.7%±3.6% and 37.5%±8.8%, respectively (Gao et al., 2018). These studies indicate the need for further improvement in achieving more desirable gene integration efficiency (similar to gene disruption efficiency of about 90%) via CRISPR-Cas9-based tools. Besides, gene insertion via HR-mediated double-strand break (DSB) repair usually demands synchronous introduction of two plasmids harboring different selection markers (Gao et al., 2018; Schwartz et al., 2017b), resulting in reduction of transformation efficiency and additional procedure for plasmids recovery.

The Cre/*lox*-site-specific recombination system derived from bacteriophage P1 is known to be appropriate for genome editing (Kawano et al., 2016; Kuhn and Torres, 2002; Richardson et al., 2017; Wu et al., 2018), and its action mechanism has been clearly described (Albert et al., 1995; Leibig et al., 2008; Yan et al., 2008). Cre recombinase can recognize and bind to *lox*P site, and can catalyze molecular recombination reactions including insertion, translocation, inversion and deletion (Albert et al., 1995). However, these reactions can retain redundant *lox*P sites in the genome, which may disturb further recombination (Albert et al., 1995). To address this issue, some mutational *lox* sites have been successfully designed (Leibig et al., 2008; Yan et al., 2008). For example, the recombination between a left element-mutant *lox* site (LE) *lox*71 and a right element-mutant *lox* site (RE) *lox*66 can generate a double element-mutant *lox* site (LE+RE) *lox*72 and a *lox*P site (Leibig et al., 2008; Yan et al., 2008). Meanwhile, *lox*P site can be deleted and *lox*72 site can be retained in the genome through rational design. In particular, the *lox*72 site, which can weakly bind to Cre recombinase, is less likely to participate in subsequent recombination (Leibig et al., 2008; Yan et al., 2008). As a result, *lox* mutants can exhibit enhanced controllability of recombination and accelerate the utilization of Cre/*lox* system in yeasts (Fickers et al., 2003; Gueldener et al., 2002; Hentges et al., 2005; Hochrein et al., 2018). Although this method can employ marker-free genome modification, it often presents low efficiency or is time-consuming. In addition, a few studies have attempted markerless gene integration via Cre/*lox* system in *Y. lipolytica*. Among them, an inspired method combining Cre/*lox* system and 26S rDNA achieved 79.2% integration efficiency in *Y. lipolytica* (Lv et al., 2019); however, the integration loci were random and the copy numbers of target genes were uncontrollable, requiring lot of time for selection and evaluation of transformants. Moreover, to rescue selection marker in the genome, an additional plasmid harboring *cre* gene must be introduced into the recombinants by another transformation, and this plasmid should subsequently be recovered. These tedious procedures require two different selection markers and further extend the experimental period.

Therefore, in the present study, a novel and effective genetic tool was designed based on Cre/*lox* system. Only a single selection marker was adequate to enable repeated, targeted, and markerless gene integration, and all unnecessary fragments could be totally excised. To demonstrate the efficacy of the developed tool, an important lipase gene from *Rhizomucor miehei* (*rml*) was chosen as a qualitative reporter, and three new plasmids were constructed. A total of seven rounds of marker-recyclable integration were implemented, and the positive colony proportions reached 90.63%–100%. The experiment conditions, feasibility, and controllability of the genetic tool were evaluated and optimized. Moreover, to confirm the universality of the developed tool, other six different genes were further integrated into *leu2* locus, and a derived tool was successfully employed to integrate another target gene into the *axp1-2* locus. The results obtained revealed that Cre/*lox*-based genetic tool could resolve the limitations of the existing methods and bring significant advancement in gene editing in yeast species.

## 2. Material and methods

### 2.1 Strains and media

*Escherichia coli* TOP10 and TOP10 F’ (Shanghai Weidi Biotechnology Co., Ltd, Shanghai, China) were applied for plasmid amplification. The plasmid hp12d-*rml* (Zhou et al., 2019) and auxotrophic *Y. lipolytica* Po1f (Madzak et al., 2000) and Po1h (Madzak et al., 2004) were respectively utilized for plasmid construction and transformation. The T4 DNA ligase, Phanta Max Master Mix polymerase, Rapid Taq Master Mix polymerase, and ClonExpress II One Step Cloning Kit were obtained from Vazyme Biotech Co., Ltd (Nanjing, China) and used for gene amplification and genome identification. The *E. coli* and *Y. lipolytica* strains were respectively cultured at 37 °C and 28 °C. All strains and plasmids used in this study are listed in Supplemental Table S1. The LB, YPD, MD, and BMSY media have been previously described (Zhou et al., 2019). The MO medium (20 mL/L oleic acid, 0.5 mL/L Tween 80, 13.4 g/L yeast nitrogen base with no amino acids, and 0.4 mg/L biotin) was derived from MD medium, and leucine (final concentration, 262 mg/L) or uracil (final concentration, 22.4 mg/L) was added for auxotrophic strains.

### 2.2 Plasmid construction

Supplemental Fig. S1 illustrates plasmid Cre-Y1 construction. The pPOX2 promoter, *cre* gene, tandem fragment of lip2t and *lox*71, and fusion segment of Upleu and *lox*71 were respectively cloned via corresponding primer pairs (Pox2-F1/Pox2-R1, Cre-F1/Cre-R1, lip2t-F1/lip2t-R1, and Upleu-F1/Upleu-R1). The four amplified products were assembled by overlap extension PCR and then double-digested with *Nde*I/*Apa*I. Subsequently, the digested product was inserted into *Nde*I/*Apa*I-digested plasmid hp12d-*rml* to construct plasmid 1296*-cre*. Furthermore, the *ura3* gene (amplification using Ura-F1/Ura-R1) was subcloned into *Nde*I-linearized 1296*-cre* by HR, generating Cre-Y1. All the plasmids were confirmed by DNA sequencing by TsingKe Biological Technology Co., Ltd. (Wuhan, China), and all the primers and *lox* sites used in this study are presented in Supplemental Table S2.

As shown in Supplemental Fig. S2, the plasmids Cre-Y1 and hp12d-*rml* were respectively double-digested using *Apa*I/*Nhe*I and *Apa*I/*Xba*I, and the two digestion products with same cohesive ends were ligated to generate the plasmid hp12d-*rml-cre*. Subsequently, the gene segment lip2t-*lox*66 was cloned by lip2t-F1/lip2t-R2 and recombined into *Avr*II/*Mlu*I-digested hp12d-*rml-cre*, forming hp12d-*rml-cre*-*lox*66. Furthermore, the fusion fragment of rc*lox*66 and partial *leu2* was amplified by leu-F2/leu-R2 and recombined with *Nde*I/*Apa*I-opened hp12d-*rml-cre*-*lox*66 to generate plasmid Cre-Y2. Similarly, the gene segment lip2t-rc*lox*71 was amplified by lip2t-F1/lip2t-R3 and used for the construction of hp12d-*rml-cre*-rc*lox*71, and hp12d-*rml-cre*-rc*lox*71 was recombined with *lox*71-partial *leu2* to obtain plasmid Cre-Y3 (Supplemental Fig. S3).

The *rml* gene of Cre-Y2 and Cre-Y3 was replaced with six different genes (*ire1, kar2, pdi, sls1, hac1*, and *vgb*) to assemble the corresponding plasmids, and their structures are presented in Supplemental Fig. S4A. Moreover, the homologous fragment Upleu, *leu2*, and target gene *rml* were respectively substituted with Upaxp, *axp1-2*, and *rol* to generate the plasmids Cre-axp1, Cre-axp2, and Cre-axp3 (Supplemental Fig. S4B). The abovementioned procedures were achieved by conventional PCR amplification, enzyme digestion, and ligation.

### 2.3 Yeast transformation, screening and induction

The integration process was performed as follows (Supplemental Fig. S5). The *Sal*I-linearized plasmid Cre-Y1 was integrated into Po1f genome based on the previously described method (Chen et al., 1997). Then, the transformants were screened twice in 5 mL of MD liquid medium supplemented with leucine (MD-Leu). Subsequently, the derivatives were inoculated into MO liquid medium supplemented with leucine and uracil (MO-Leu-Ura) for inducible expression of *cre* gene. To isolate single colony, the cell suspension solution was streaked onto YPD plate, and the selected colonies were respectively transferred onto YPD and MD-Leu plates for phenotype verification, where the positive colonies could only grow on YPD medium. In addition, the genomic DNA of positive colonies was extracted for further identification via PCR amplification with appropriate primers and DNA sequencing by TsingKe Biological Technology Co., Ltd. (Wuhan, China). Thus, the initial strain FY1 was obtained. The subsequent integration was almost similar to the first-round integration; however, Cre-Y2 or Cre-Y3 was linearized by *Apa*I, leucine was not required in MD or MO medium, and YPD medium was replaced with BMSY medium containing 10 mL/L glyceryl tributyrate to select colonies with high RML activity.

### 2.4 Shaking flask culture and lipase activity determination

Transformants harboring clear and large transparent halos were inoculated into 5 mL of YPD for 24 h. Then, the cultures were transferred into 500-mL shake flasks containing 30 mL of BMSY medium and incubated for 120 h to produce RML. The RML activity in cell culture supernatant was evaluated using previously described method (Zhou et al., 2019). All the experiments were performed in triplicate, and the results were analyzed by ANOVA with Duncan’s multiple range test at *P* < 0.05.

### 2.4 Determination of gene copy number and transcriptional analysis

Based on the protocol of ChanQ™ Universal SYBR qPCR Master Mix (Vazyme Biotech Co., Ltd, Nanjing, China), the *rml* gene copy numbers were determined by real-time qPCR performed on StepOnePlus apparatus with StepOne software version 2.3 (Applied Biosystems, Foster City, CA, USA). The endogenous *act1* was set as reference gene, and the plasmids pMD19-*act1* and hp4d-*rml* (Zhou et al., 2019) were utilized for establishing standard curves, respectively. The *rml* gene transcriptional level analysis was conducted by TsingKe Biological Technology Co., Ltd (Wuhan, China). Each sample was collected in triplicate.

## 3. Results

### 3.1 Genetic tool design and integration process analysis

The novel Cre/*lox*-based genetic tool comprised three plasmids, namely, Cre-Y1, Cre-Y2, and Cre-Y3. Plasmid Cre-Y1 was applied for generating the initial strain, and plasmids Cre-Y2 and Cre-Y3 were employed to iteratively introduce target genes into the host strain. Cre-Y1 consisted of a yeast selection marker, *cre* gene expression cassette, bacterial element (antibiotic resistance gene and origin of replication), homologous fragment of target locus in the genome (HF-A), and two convergently oriented *lox* mutants. The *cre* expression cassette should adopt an inducible promoter to control the production of Cre recombinase. Cre-Y2 was derived from Cre-Y1 by replacing HF-A and two convergently oriented *lox* mutants with another homologous fragment of target locus (HF-B), a *lox* mutant and its reverse complement (rc*lox*), and a target gene expression cassette. To avoid excising original segments from the genome, HF-B should be close to HF-A. Cre-Y3 was similar to Cre-Y2, but with the corresponding *lox* mutants of Cre-Y2 (e.g., LE for Cre-Y2 and RE for Cre-Y3). All of the tool elements were flexible and nonspecific; thus, the Cre/*lox*-based genetic tool was appropriate for different species.

In the present study, the plasmid elements used for *Y. lipolytica* were as follows: the auxotroph Ura3 (UniProtKB, Q12724) was adopted as selection marker, *cre* expression cassette was assembled as pPOX2-*cre*-lip2t (pPOX2 promoter, *cre* gene, and *lip2* terminator, GenBank Accession Nos. AJ001300.1, AB449974.1, and AJ012632.1), bacterial element (Amp-ori) and target gene expression cassette (*rml*) were subcloned from plasmid hp12d-*rml*, and HF-B and HF-A were respectively designed as *leu2* gene (GenBank Accession No. AF260230.1) and its upstream segment (Upleu) for the convenience of plasmid construction and linearization. In addition, two *lox*71 sites, a *lox*66 site and a reverse complement of *lox*66 site (rc*lox*66) and a reverse complement of *lox*71 site (rc*lox*71) and a *lox*71 site were used in plasmids Cre-Y1, Cre-Y2, and Cre-Y3, respectively.

To demonstrate repeated and markerless gene integration in *Y. lipolytica* via Cre/*lox*-based genetic tool, the experimental process was as follows. First, Cre-Y1 containing two *lox*71 sites was introduced into the Upleu locus of Po1f to construct an initial strain harboring a single *lox*71 site. Under the condition of suitable screening and induction, recombination between two *lox*71 sites could remove unnecessary plasmid elements and produce the initial strain. Then, Cre-Y2 was integrated into *leu2* locus of the initial strain, forming a transitional strain harboring three *lox* sites (*lox*71, *lox*66, and rc*lox*66) in the genome. Various recombination reactions among these three *lox* sites could excise unnecessary DNA fragments and retain a *lox*72 site, an rc*lox*66 site, and a target gene expression cassette in the genome. Meanwhile, two types of stable strains were eventually generated. Similarly, integration of Cre-Y3 produced strains harboring two target gene expression cassettes. Consequently, repeated, targeted, and markerless gene integration could be efficiently realized following iterative integration of Cre-Y2 and Cre-Y3.

### 3.2 First-round integration to construct initial strain FY1

To construct strain harboring a single *lox*71 fragment, the plasmid Cre-Y1 was inserted into auxotrophic host Po1f (Fig. 1). The transformed cell pellet was screened twice in 5 mL of MD-Leu medium to enrich the Ura^+^ recombinant strain Po1f/Cre-Y1 (Ura^+^ and Leu^-^) and avoid interference from untransformed Po1f cells (Supplemental Fig. S5). Subsequently, the culture solution containing Po1f/Cre-Y1 was transferred into MO-Leu-Ura medium for 12 h with oleic acid as the sole carbon source, which could activate pPOX2 to induce the expression of *cre* gene. In this case, the produced Cre recombinase could mediate deletion reaction between two repeat *lox*71 sites in the Po1f/Cre-Y1 genome, causing excision of the bacterial element (Amp-ori), selection marker (Ura3), and *cre* expression cassette, further generating Ura^-^ recombinants. Meanwhile, the extra added uracil allowed Ura^-^ recombinants to grow normally. The induced cell suspension was streaked on YPD plate for isolating single colony with Ura^-^ genotype, and the phenotype of the isolated colonies were determined on YPD and MD-Leu media. As shown in Supplemental Fig. S6A, a total of 32 recombinants were randomly chosen and all of them could not grow on MD-Leu plate, proving that the selection marker Ura3 was recycled successfully. Moreover, 10 positive colonies were picked up randomly and identified by genome PCR (Supplemental Fig. S7A), and the PCR product obtained via primers Upleu-F3/Upleu-R3 was further sequenced. The results shown in Supplemental Fig. S8A indicated that only a single *lox*71 site was retained in Upleu locus of the eventually obtained strain (named FY1). This finding also revealed that all recombination and integration events occurred as expected (Fig. 1), and the adopted plasmid elements of the novel genetic tool were well suitable for *Y. lipolytica*.

**Fig. 1.**
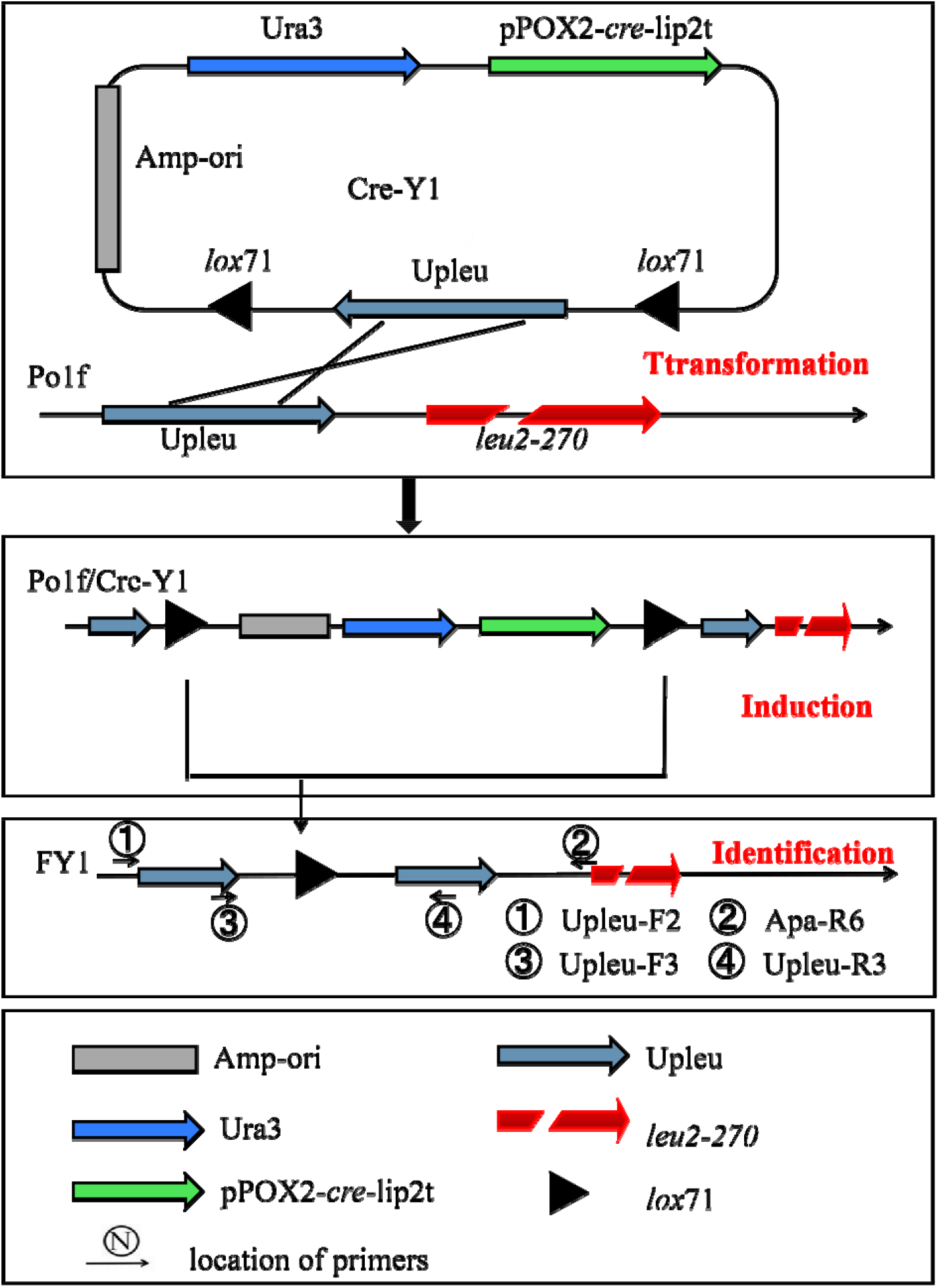
Schematic diagram of first-round integration to obtain initial strain FY1. The Cre-Y1 was linearized at homologous fragment Upleu and introduced into Po1f. The Po1f/Cre-Y1 was screened twice in MD-Leu medium and then inoculated into MO-Leu-Ura medium for inducible expression of *cre* gene. The bacterial element (Amp-ori), selection marker (Ura3) and *cre* expression cassette (pPOX2-*cre*-lip2t) were removed by deletion reaction of two *lox*71 sites, retaining a single *lox*71 site in the FY1 genome. Primers used for PCR amplification are labeled in this figure and described in Supplemental Table S2.

### 3.3 Second-round integration and analysis

To evaluate whether the designed plasmids could realize markerless gene integration, Cre-Y2 was linearized within *leu2* and introduced into FY1, to obtain the strain FY1/Cre-Y2. As shown in Fig. 2, three *lox* sites (*lox*71, *lox*66, and rc*lox*66) were noted in the FY1/Cre-Y2 genome, and three kinds of recombination reactions could occur when Cre recombinase was expressed. In the first-type recombination (a), deletion reaction occurred between the *lox*71 and *lox*66 sites in the FY1/Cre-Y2 genome, and the strain FY2-*rml* carrying *rml* expression cassette was obtained. In the second-type recombination (b), the *lox*71 and rc*lox*66 sites were combined with Cre proteins, producing a *lox*72 site and reverse complement of *lox*P site (rc*lox*P). Here, the *lox*72 site was hardly bound to Cre recombinase, and the rc*lox*P site could not stably coexist with the unreacted rc*lox*66 site in the genome, which could recombine with Cre recombinase again, generating FY2-rc*rml*, a derived strain carrying the reverse complement of *rml* gene expression cassette (rc*rml*). In the third-type recombination (c), the *lox*66 and rc*lox*66 sites were recombined for inversion reaction, forming a transitional strain that also contained three *lox* sites similar to those of FY1/Cre-Y2. The recombination reaction among the three *lox* sites continued until the first or second type of recombination occurred, and the strain FY2-*rml* or FY2-rc*rml* was finally generated. Consequently, two kinds of stable strains were obtained from three types of recombination reactions in second-round integration. Here, all of the unnecessary and redundant fragments, including Upleu, *leu2-270*, bacterial element, selection marker, and *cre* expression cassette, were completely deleted, and only gene expression cassette and rc*lox*66 site remained in the genome.

**Fig. 2.**
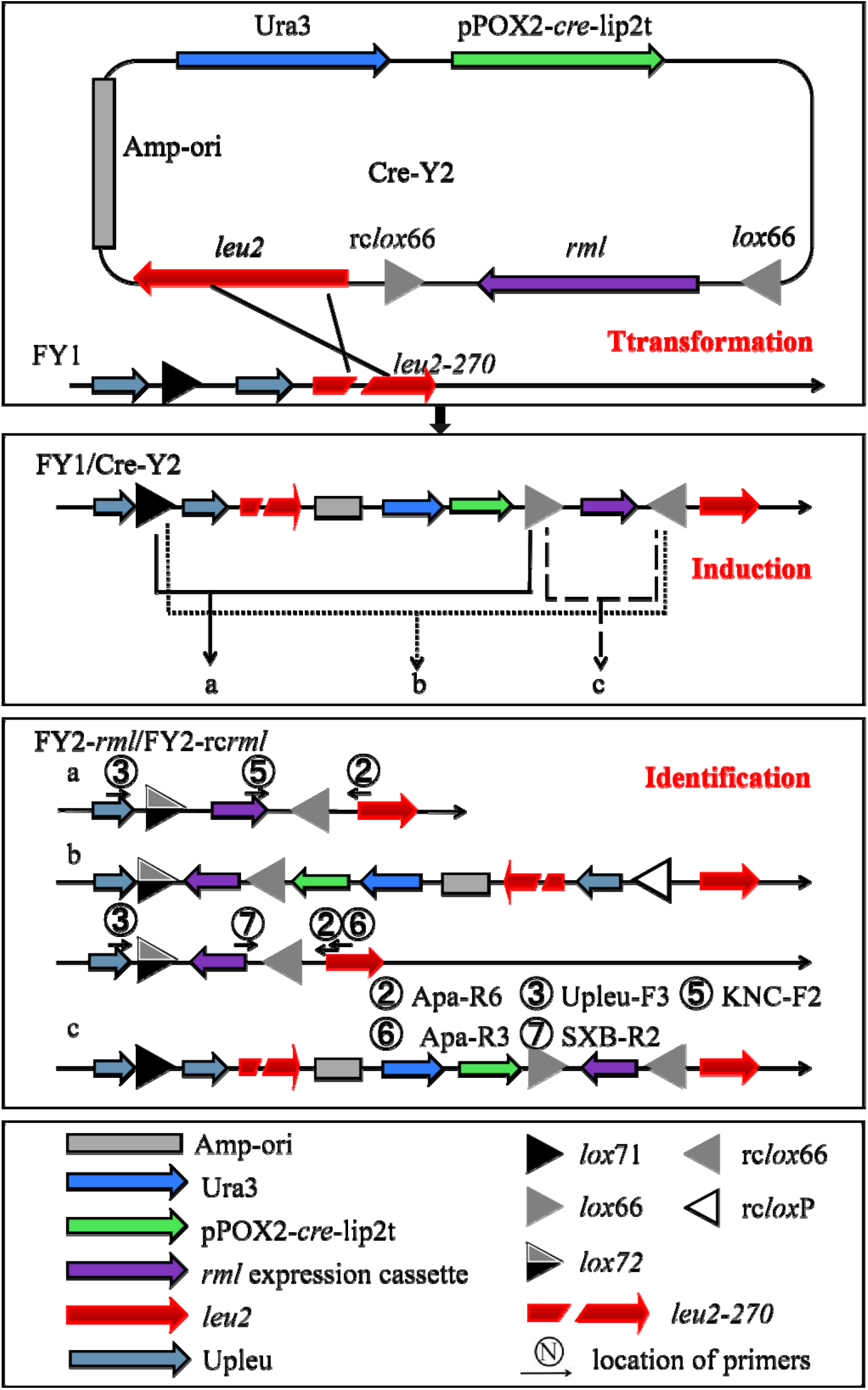
Schematic diagram of second-round integration to generate FY2-*rml*/FY2-rc*rml*. The Cre-Y2 was linearized on *leu2* and integrated into the FY1 genome to obtain recombinant strain FY1/Cre-Y2. The FY1/Cre-Y2 was screened twice in MD medium and then induced in MO-Ura medium for the production of Cre protein. Therein, three kinds of different recombination reactions for *lox* sites were noted in the genome: deletion reaction between *lox*71 and *lox*66 sites (a), inversion reaction between *lox*71 and rc*lox*66 sites (b), and inversion reaction between *lox*66 and rc*lox*66 sites (c). Eventually, two forms of stable strains FY2-*rml* and FY2-rc*rml* were produced. Meanwhile, all of the unnecessary and redundant sequences were excised, the *rml* expression cassette was integrated into the genome, and the remaining rc*lox*66 site locus was employed in third-round integration. Primers used for PCR amplification are labeled in this figure and described in Supplemental Table S2.

### 3.4 Induction time optimization in second-round integration

In the present study, oleic acid induced pPOX2 promoter controlled the production of Cre recombinase in MO liquid medium. Therein, induction time affected Cre protein concentration in the cell, and further influenced the recombination efficiency in the engineered strains. In the first-round gene integration, there were two *lox* sites in the yeast genome. The recombinants were induced for sufficiently long time to obtain high proportion of positive colonies. However, in subsequent gene integrations, there were three or more *lox* sites in the genome and the recombination reaction was complex and diverse. As reported in previous studies (Albert et al., 1995; Suzuki et al., 2005), high concentration Cre protein is more likely to mediate recombination in *lox*72 site, resulting in the removal of the target gene expression cassette from the engineering strain genome. Therefore, induction of *cre* gene expression should be controlled in an appropriate time range. In the present study, to determine the optimal induction time, the recombinant strain FY1/Cre-Y2 was incubated in MO-Ura liquid medium for 4–16 h and streaked onto YPD medium every 2 h (approximately one cell generation cycle). Furthermore, 32 recombinants were randomly selected from each YPD medium and then transferred onto MD and BMSY media. The proportions of positive colonies under different induction times were examined, and the results are shown in Table 1 and Supplemental Fig. S6B. All the selected recombinants could complete recombination reaction after 12 h of induction. For more accurate confirmation, 64 colonies induced for 12 h were chosen and none of them could normally grow on MD medium. Consequently, the induction time was set as 12 h, which could be further increased if the proportion of positive colonies decreased in multi-round integrations.

**Table 1.**
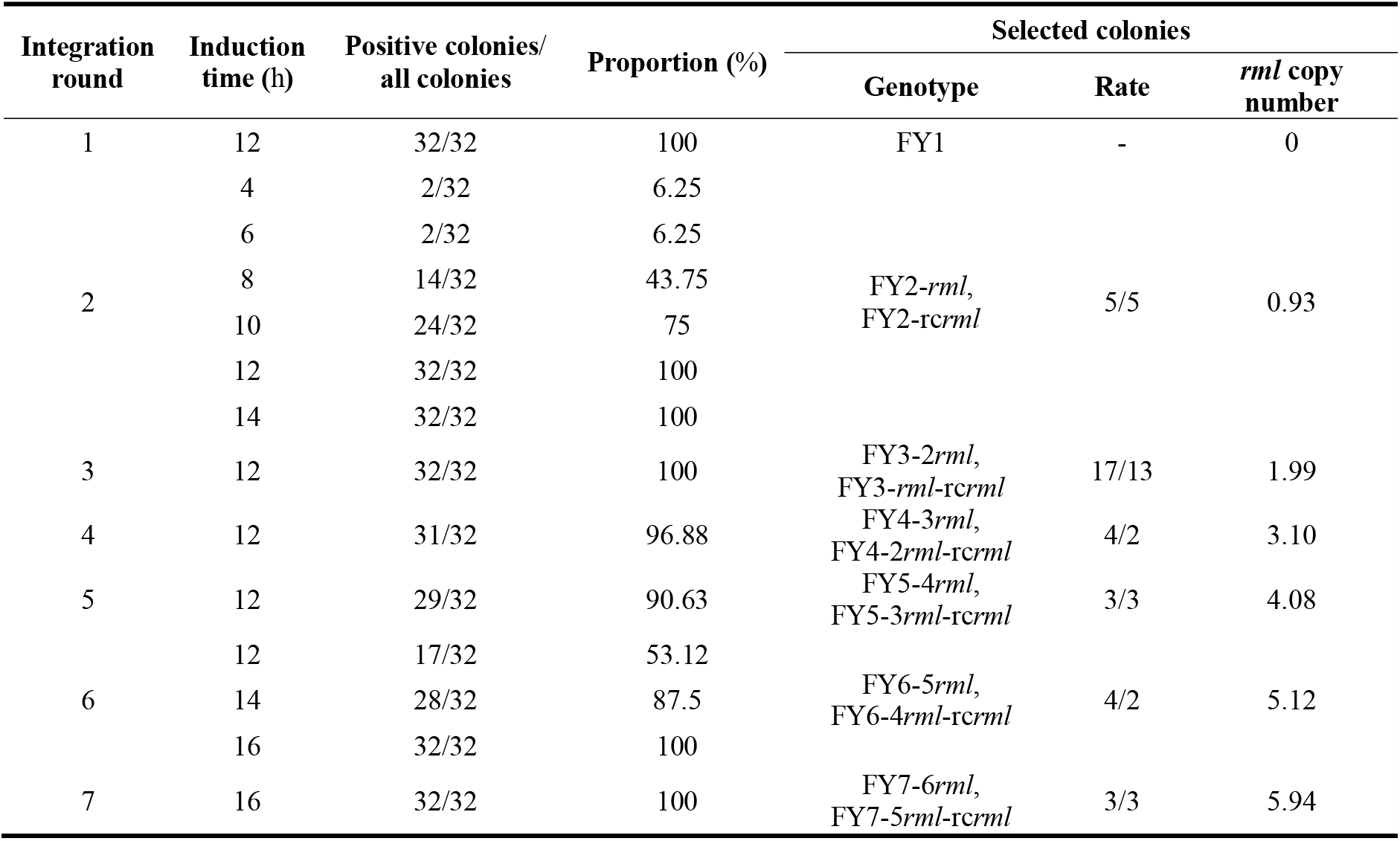
**Positive colony proportion, genotype rate, and *rml* copy number determined in this study.**

A total of 10 positive colonies were randomly chosen for identification via PCR sequencing (Supplemental Fig. S7B). The *rml*/rc*rml* expression cassette and rc*lox*66 site in all the selected colonies matched with the genome of FY2-*rml*/FY2-rc*rml* (Supplemental Figs. S8B, C), demonstrating that the second-round gene integration was performed according to the predictive analysis (Fig. 2). In addition, for the convenience of subsequent protein production evaluation, strains that do not carry rc*rml* expression cassette were employed for further gene integration, because the existing plasmids and method (Zhou et al., 2019) are more applicable to construct strains only harboring *rml* expression cassettes.

### 3.5 Iterative insertion of Cre-Y3 and Cre-Y2

To achieve third-round integration, Cre-Y3 was utilized to transform FY2, and the schematic diagram is presented in Fig. 3. Similar to the second-round integration, there were three *lox* mutants (rc*lox*66, rc*lox*71, and *lox*71) and a *lox*72 site in FY2-*rml*/Cre-Y3 genome, resulting in three different recombination reactions and eventually generating two genotypes of the recombinant strains (FY3-2*rml* and FY3-*rml*-rc*rml*). Subsequently, FY3-2*rml* and FY3-*rml*-rc*rml* were distinguished by PCR amplification with primer pairs KNC-F2/Apa-R6 and SXB-R2/Apa-R3 (Supplemental Fig. S7C). The obtained products were further sequenced and the results (Supplemental Figs. S8D, E) were consistent with the prediction presented in Fig. 3. Thus, another copy of *rml* gene was targeted on *leu2* locus and all unnecessary DNA fragments were excised in the third-round integration.

**Fig. 3.**
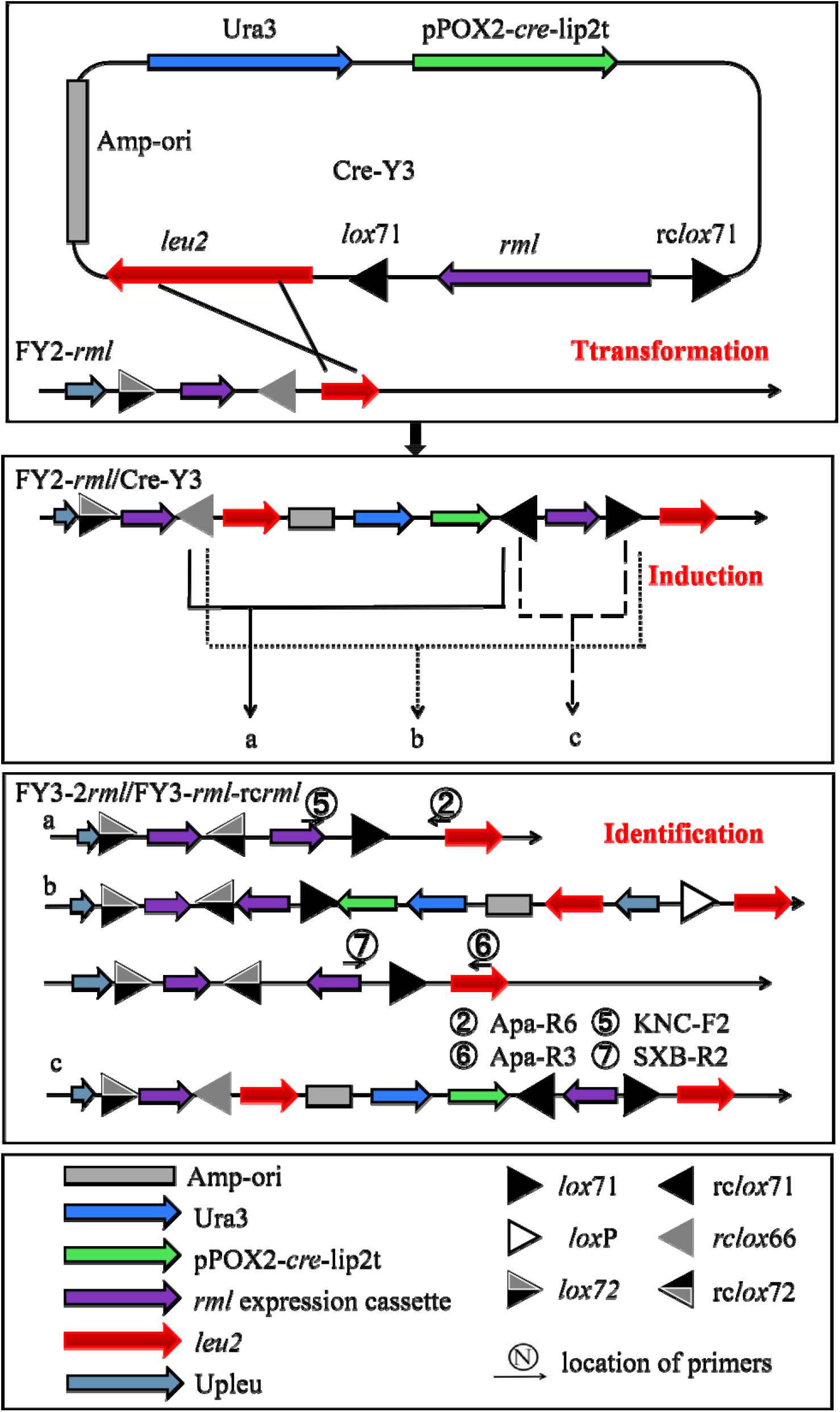
Schematic diagram of third-round integration to construct FY3-2*rml*/FY3-*rml*-rcr*ml*. The ApaI-linearized Cre-Y3 was used to transform FY2-*rml* 3#, generating the strain FY2-*rml*/Cre-Y3. There were three kinds of different recombination reactions for *lox* sites in FY2-*rml*/Cre-Y3 genome. Herein, the recombination reactions excised unnecessary DNA fragments (a *leu2*, Amp-ori, Ura3, and pPOX2-*cre*-lip2t) and eventually generated two kinds of stable strains FY3-2*rml* and FY3-*rml*-rc*rml*. The remaining *lox*71 site could be used for subsequent integration process. Primers used for PCR amplification are labeled in this figure and described in Supplemental Table S2.

**Fig. 4.**
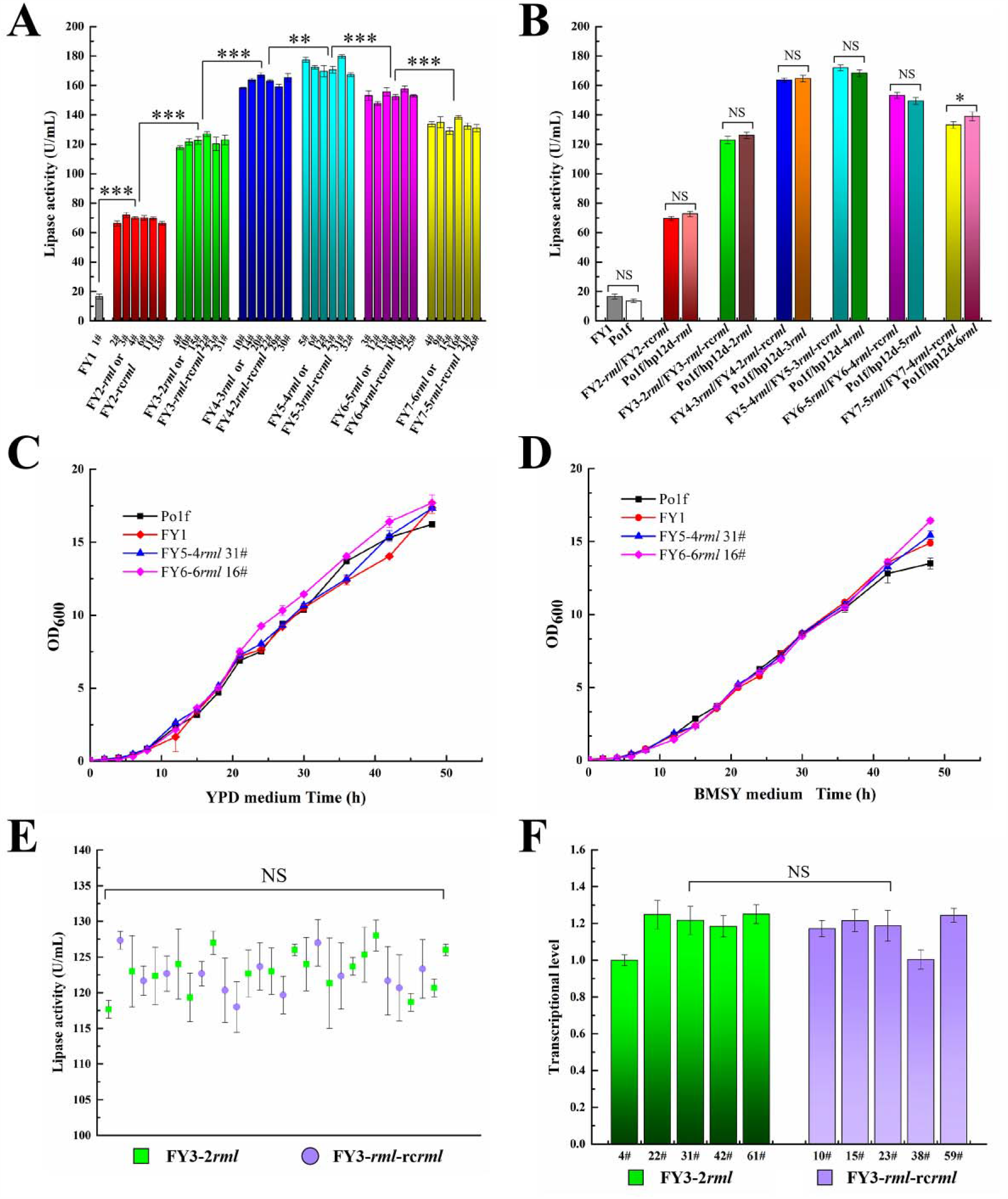
(A) Lipase activity of recombinant strains harboring different copies of *rml* expression cassettes. (B) Comparison of lipase activity of recombinant strains obtained via different methods. Data are the mean values of each kind of recombinant strain. The strains Po1f and FY1 showed low lipase activity owing to expression of autologous lipase genes. Weakly significant difference was observed between the strains FY7-6*rml*/FY7-5*rml*-rc*rml* and Po1f/hp12d-6*rml* (*P=*0.044). (C) Lipase activity of transformants from third-round integration. (D) Transcriptional level analysis of *rml* mRNA from FY3-2*rml* and FY3-*rml*-*rcrml*. The endogenous *act1* was adopted as reference gene, and the *rml* transcriptional level of FY3-2*rml* 4# was defined as 1. (E) Growth curves of Po1f, FY1, FY5-4*rml* 31#, and FY7-6*rml* 13# in YPD medium. (F) Growth curves of Po1f, FY1, FY5-4*rml* 31#, and FY7-6*rml* 13# in BMSY medium. NS, ^*, **^, and ^***^indicate non-significant, *P* < 0.05, *P* < 0.01, and *P* < 0.001, respectively.

Similarly, Cre-Y2 was introduced into FY3-2*rml* to generate FY4-3*rml*/FY4-2*rml-* rc*rml*, and Cre-Y3 was used to transform FY4-2*rml* for constructing FY5-4*rml*/FY5-3*rml-*rc*rml* (Supplemental Fig. S9A). Subsequently, Cre-Y2 and Cre-Y3 were iteratively integrated into the corresponding recombinant strains to obtain FY6-5*rml*/FY6-4*rml-* rc*rml* and FY7-6*rml*/FY7-5*rml-*rc*rml*. The genome PCR identification results (Supplemental Fig. S9B) proved that repeated, targeted, and markerless gene integration was successfully implemented by the novel genetic tool.

### 3.6 Increasing induction time in sixth and seventh rounds of integration

As shown in Table 1, the positive colony proportions slightly decreased in multi-round integration processes, which might be owing to the following possibility: an additional *lox*72 site could have been inserted into the genome in each round of gene integration, except in the first-round integration, and all of the *lox* sites can bind to Cre recombinase (Albert et al., 1995; Leibig et al., 2008; Yan et al., 2008). Thus, *lox*72 site might disturb the combination between LE/RE *lox* sites and Cre recombinase, resulting in increasingly less recombination reactions during the induction process and a decrease in efficiency in multi-round integrations. Nevertheless, an increase in induction time caused higher accumulation of Cre protein in the cell, and more Cre recombinase could recognize LE/RE mutants and mediate effective recombination.

Accordingly, the transformants were induced for longer time to obtain higher recombination efficiency in the sixth-round integration. As a result, the proportion of positive colonies was enhanced, and 16 h was adequate to complete recombination in the engineered strains (Table 1 and Supplemental Fig. S6C). In the seventh-round integration, 16 h was employed as the induction time and all the chosen colonies could not grow on MD medium (Supplemental Fig. S6C). Therefore, recombination efficiency might be roughly inversely proportional to the integration round; however, this problem could be alleviated by increasing the induction time in MO-Ura liquid medium. Finally, seven rounds of gene integration were implemented in this study, and the positive colony proportions reached 90.63%–100% with an induction time of 12–16 h (Table 1). In fact, the realized integration rounds were adequate to meet the demands of engineering strains construction, and the potential of the developed genetic tool could also be explored in future research.

### 3.7 Protein expression level and growth characteristics evaluation

The recombinants harboring different *rml* gene copies were cultured in BMSY liquid medium for protein production, and the results are presented in Fig. 5A. The recombinant strains FY5-4*rml* 31# showed the highest RML expression level, while the lipase activity of FY6-5*rml*/FY6-4*rml*-rc*rml* and FY7-6*rml*/FY7-5*rml*-rc*rml* was slightly lower. Thus, gene integration was terminated in the seventh round. Moreover, several colonies were chosen as representatives for each type of the recombinant strains, and their *rml* gene copies were determined via qPCR. As shown in Table 1, all the transformants carried the same gene copy number as previously predicted, indicating that no *rml* expression cassette was excised from the genome and also confirming the excellent controllability and stability of the genetic tool.

To evaluate the protein expression level of the recombinant strains generated via the developed genetic tool, the plasmids hp12d-5*rml* and hp12d-6*rml* were constructed according to the previous method (Zhou et al., 2019). Subsequently, the plasmid hp12d-n*rml* (n=1–6) were linearized and introduced into Po1f to form the recombinant strains Po1f/hp12d-n*rml* (carrying n copies of *rml* expression cassette, n=1–6). Then, Po1f/hp12d-n*rml* (n=1–6) and engineered strains from seven rounds of gene integration were respectively inoculated into BMSY medium under the same culture conditions, and their lipase activity was measured (Fig. 5B), which revealed that the two types of recombinant strains showed similar protein expression level. Therefore, it can be concluded that genome editing via the developed genetic tool did not affect target protein production of the engineering strains. When compared with the previous method (Zhou et al., 2019), the developed genetic tool has many advantages such as utilization of reusable marker, only requiring three plasmids, no gene redundancy, and flexible direction of gene expression cassette. Moreover, the novel genetic tool could assemble engineering strains harboring any copies of the target gene, only requiring increase/decrease in integration round.

Furthermore, the strains Po1f, FY1, FY5-4*rml* 31#, and FY7-6*rml* 13# were cultivated in YPD and BMSY media, respectively, and their growth characteristics were tested by measuring the optical density at 600 nm (OD_600_) at different incubation times (Figs. 5C, D). The results revealed no observable differences in the growth curves of the four types of strains, indicating that the genome modification in *leu2* locus via Cre/*lox*-based genetic tool did not influence cell growth.

### 3.8 Comparison of two-genotype recombinant strains

Two genotypes of the recombinant strains were generated in each round of integration (except for the first round), and their numbers were presented in Table 1. The rates of two forms of strains were approximately equal, suggesting that there was nearly equal possibility to choose a strain with certain genotype. The colonies with high lipase activities were selected for protein production evaluation. The results shown in Table 1 also implied that the protein expression levels of the recombinant strains in each round of integration were similar, which was confirmed in Fig. 5A. In contrast, some previous studies have reported that the relative arrangement of multiple genes might affect corresponding transcription and protein production in host cells (Eszterhas et al., 2002; Lee et al., 2014). Thus, a total of 30 positive colonies from the third-round integration were identified via the primer pairs KNC-F2/Apa-R6 and SXB-R2/Apa-R3 (data not shown), and their lipase activities were measured. The results presented in Fig. 5E showed that the RML activity of each recombinant randomly changed within a proper range. Moreover, transcriptional level analysis further indicated that there was no significant difference in the transcriptional quantity of *rml* mRNA isolated from FY3-2*rml* and FY3-*rml*-rc*rml* cells (Fig. 5F). Thus, the relative direction of *rml* expression cassettes had no observable effect on protein production in this study, which might be owing to the difference in the host type. Accordingly, both the genotypes of the strains were concluded to be suitable for RML production in *Y. lipolytica*.

### 3.9 Successfully expanding to six different genes and axp1-2 locus

To evaluate the universality of the novel genetic tool, six genes with varied lengths and resources were additionally integrated into the engineered strain FY5-4*rml*. The genes *ire1, kar2, pdi, sls1*, and *hac1* are the homologous genes of *Y. lipolytica*, and *vgb* is the coding gene of *Vitreoscilla* hemoglobin. These genes may differentially affect RML expression level, which determines their integration times. The corresponding plasmids were derived from Cre-Y2 and Cre-Y3 by replacing the *rml* gene with different target genes (Supplemental Fig. S4A), and then used to transform FY5-4*rml*. The subsequent integration processes were similar to those of Cre-Y2 and Cre-Y3, and the proportions of positive colonies were all above 90% (Table 2), confirming the availability of the developed genetic tool for repeated, targeted, and markerless integration of different genes.

**Table 2.**
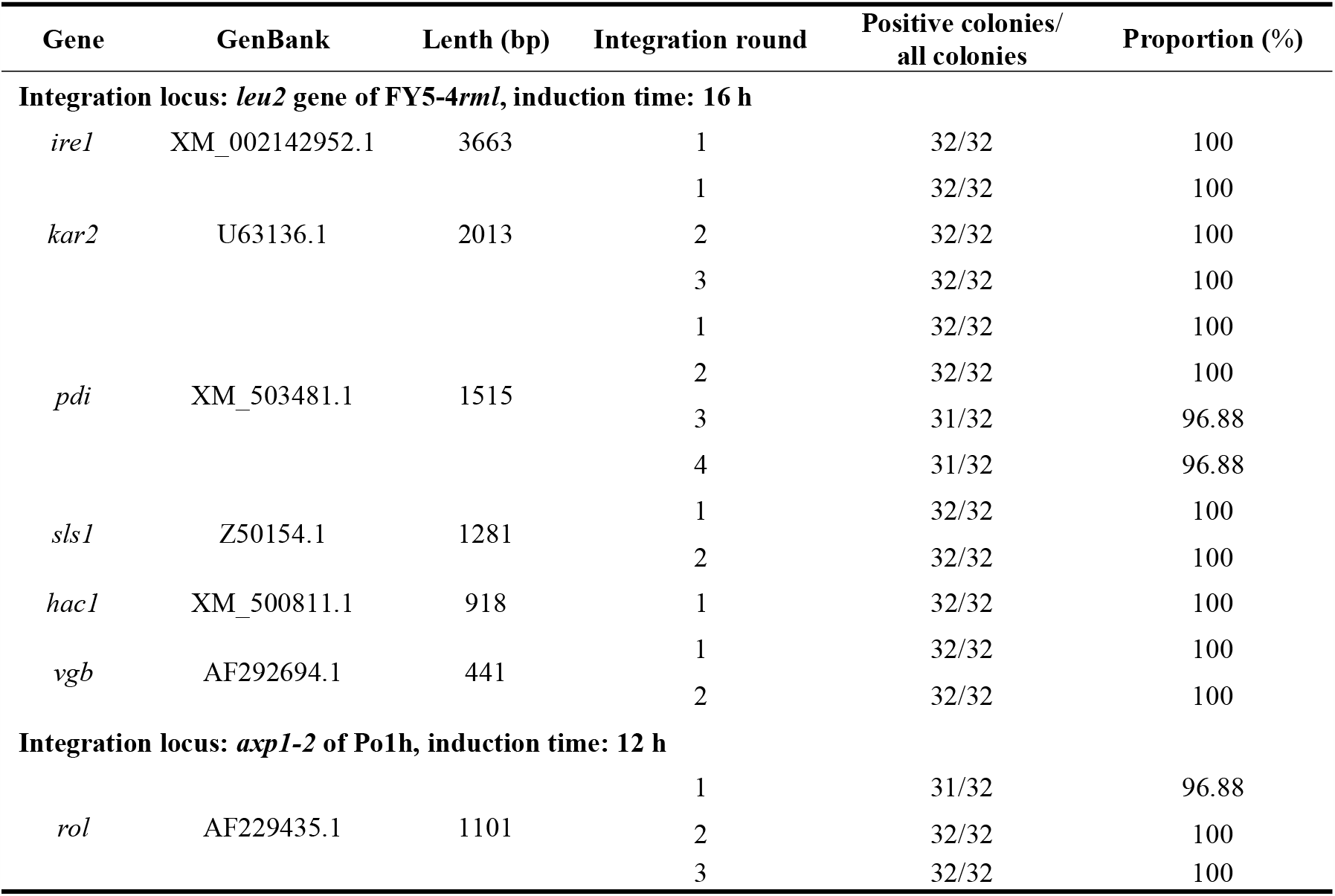
Integration rates of various genes, loci and strains in this study.

Furthermore, a derived tool was established to verify whether the developed genetic tool could be applied for other target genes, integration loci, and strains. For this purpose, three plasmids containing *Rhizopus oryzae* lipase (*rol*) gene were respectively constructed based on Cre-Y1, Cre-Y2, and Cre-Y3 (Supplemental Fig. S4B), and integrated into defective *axp* gene locus (*axp1-2*) of *Y. lipolytica* Po1h. The transformation, screening and induction, isolation of positive colonies, and identification processes were similar to those employed for *rml* gene integration into *leu2* locus of Po1f. First, the Cre-axp1 plasmid was linearized by *Nsi*I and inserted into the Upaxp locus to generate an initial strain. Then, the Cre-axp2 and Cre-axp3 plasmids were digested with *Cla*I and iteratively introduced into the *axp1-2* locus for repeated, targeted, and markerless integration of *rol* in Po1h. As a result, three rounds of integration were conducted and all of the integration rates were above 96%. These findings revealed that such extension of the novel genetic tool is feasible and convenient.

## 4. Discussion

In the present study, a Cre/*lox*-based genetic tool was precisely and skillfully designed. The plasmids Cre-Y2 and Cre-Y3 adopted two inversely oriented *lox* sites, and the Ura3 marker cannot be removed from the genome if the strain does not undergo integration process as expected. Thus, the Ura^+^ strain could be distinguished by isolating positive colonies on the MD medium. It must be noted that except for the *cre* gene and *lox* sites, all of the genetic tool elements are nonspecific and replaceable. Any DNA sequence can be chosen as the homologous fragment as long as no gene is disrupted, and the length of the homologous fragment is very flexible because of the excision of unnecessary sequences. The *rml* gene in plasmids Cre-Y2 and Cre-Y3 can be replaced by heterologous and endogenous genes of *Y. lipolytica* to accomplish repeated markerless integration, and the plasmid construction process is simple and convenient. In addition, the Cre/*lox*-based genetic tool can also be applied to other genes and loci if the corresponding plasmid elements are available.

Recently, several genetic tools have been developed for markerless gene integration of *Y. lipolytica* based on CRISPR-Cas9 system, Cre/*lox* system and Ura/5-fluoroorotic acid resistance method (Supplemental Fig. S10) (Gao et al., 2018; Holkenbrink et al., 2018; Lv et al., 2019; Schwartz et al., 2017b). As shown in Supplemental Table S3, the integration efficiency markedly fluctuates with different genome loci and single-guide RNAs (sgRNAs) for CRISPR-Cas9-mediated gene integrations (Holkenbrink et al., 2018; Schwartz et al., 2017b). According to most of the existing Cre/*lox*-based methods (Holkenbrink et al., 2018; Lv et al., 2019), the DNA fragment can be integrated into the host genome via gene insertion or gene replacement. However, insertion event may introduce unnecessary DNA segments into the genome, whereas replacement events, which occur occasionally, can avoid this issue, but requires substantial screening of target transformants. Furthermore, while the Ura/5-fluoroorotic acid resistance method (Ura pop-in/pop-out method) can also accomplish markerless gene integration, reverse mutation is difficult to avoid in the marker recovery process. In addition, new plasmids carrying different sgRNA or homologous fragments must be constructed to repeatedly realize gene integration in a specific genome locus via the three above-mentioned genetic tools. These limitations can be completely overcome by the novel genetic tool designed in the present study. The target gene is integrated into the host genome via efficient, targeted, and repeatable gene insertion event, and the introduced non-target sequences can be deleted by recombination reaction of the *lox* sites. Moreover, despite the occurrence of inefficient insertion event in the integration process, a few recombinants can be enriched via twice auxotroph screening. The existing methods based on CRISPR-Cas9 and Cre/*lox* systems require two different selection markers and additional plasmid recovery procedure (Gao et al., 2018; Holkenbrink et al., 2018; Lv et al., 2019; Schwartz et al., 2017b). In contrast, only a single marker is required for the novel Cre/*lox*-based genetic tool, and the process of gene integration, markers reuse, and plasmid recovery can be combined and accomplished simultaneously, which can significantly reduce the experimental period and workload. Overall, the novel genetic tool developed in this study can overcome many limitations such as low HR rate, limited selection marker, random integration site, and additional plasmid recovery.

In conclusion, a powerful and efficient Cre/*lox*-based genetic tool for repeated, targeted, and markerless gene integration was developed in this study. In practice, only a single selection marker can enable iterative integration of target gene into the genome, which can facilitate genetic modifications of the host strains. Furthermore, in each round of integration, all of the unnecessary segments are removed, the relative direction of each target gene expression cassette is controllable, and the gene recombination on genome is foreseeable. These characteristics can significantly decrease gene redundancy in the engineering strain, making it convenient for the use of host strain in pharmaceutical, food, and other health-related areas. Besides, the developed genetic tool is also suitable for various other genes and loci. The findings of this study significantly broaden the application of Cre/*lox* system and strengthen the capabilities for genome modification in *Y. lipolytica*.

## Acknowledgments

We acknowledge financial support by the National Natural Science Foundation of China (No: 41972319), the National High Technology Research and Development Program of China (Nos: 2013AA065805 and 2014AA093510), and the Fundamental Research Funds for HUST (Nos. 2017KFXKJC010 and 2017KFTSZZ001). We appreciate the Research Core Facilities for Life Science (Huazhong University of Science and Technology) for their valuable assistance in qPCR measurements. We thank International Science Editing (http://www.internationalscienceediting.com) for editing this manuscript.

## Author contributions

Q.Z. and L.J. conducted the project design. Q.Z., L.J., W.L. and Y.L. carried out the experiments. Q.Z., L.J., Z.H. and H.Z. helped with the formal analysis. Q.Z., L.J., L.X. and Y.Y. conducted the manuscript preparation.

