## supplemental file for "A novel and effective Cre/*lox*-based genetic tool for repeated, targeted and markerless gene integration"

**This document includes:**

Supplemental Tables S1 to S3

Supplemental Figures S1 to S10

**Supplemental Table S1. Strains and plasmids used in this study.**

| **Plasmid/Strain** | **Description** | **Source** |
| --- | --- | --- |
| **Plasmid** |  |  |
| T*-cre* | pMD19T derivative, carrying *cre* gene, stored in our laboratory |  |
| hp12d-n*rml*(n=1–4) | expression vector carrying n (n=1–4) copies of *rml* expression cassettes | (Zhou et al., 2019) |
| hp12d-5*rml* | expression vector carrying 5 copies of *rml* expression cassettes | This study |
| hp12d-6*rml* | expression vector carrying 6 copies of *rml* expression cassettes | This study |
| 1296*-cre* | hp12d-*rml* derivative, carrying *cre* expression cassette and two *lox*71 sites | This study |
| Cre-Y1 | 1296*-cre* derivative, carrying marker Ura3 | This study |
| hp12d-*rml-cre* | Cre-Y1 and hp12d-*rml* derivative, carrying *rml* expression cassette and a *lox*71 site | This study |
| hp12d-*rml-cre*-*lox*66 | hp12d-*rml-cre* derivative, carrying a *lox*66 site | This study |
| Cre-Y2 | hp12d-*rml-cre*-*lox*66 derivative, carrying a *lox*66 and a rc*lox*66 | This study |
| hp12d-*rml-cre*-rc*lox*71 | hp12d-*rml-cre* derivative, carrying a rc*lox*71 | This study |
| Cre-Y3 | hp12d-*rml-cre*- rc*lox*71 derivative, carrying a rc*lox*71 and a *lox*71 | This study |
| pMD19-*act1* | pMD19T derivative, carrying *act1* gene | (Zhou et al., 2019) |
| **Strain** |  |  |
| W29 | wild-type *Y. lipolytica* strain |  |
| Po1f | *MatA*, *leu2-270*, *ura3-302*, *xpr2-322*, *axp1-2*, Leu^-^, Ura^-^, △AEP, △AXP | (Madzak et al., 2000) |
| Po1h | *MatA*, *ura3-302*, *xpr2-322*, *axp1-2*, Ura^-^, △AEP, △AXP | (Madzak et al., 2004) |
| Po1f/Cre-Y1 | Po1f derivative, carrying Cre-Y1 | This study |
| FY1 | Po1f derivative, carrying a *lox*71 site at *leu2* gene locus | This study |
| FY1/Cre-Y2 | FY1 derivative, carrying Cre-Y2 | This study |
| FY2-*rml* | FY1 derivative, carrying a *rml* expression cassette | This study |
| FY2-rc*rml* | FY1 derivative, carrying a rc*rml* expression cassette | This study |
| FY2-*rml*/Cre-Y3 | FY2-*rml* derivative, carrying Cre-Y3 | This study |
| FY3-2*rml* | FY2-*rml* derivative, carrying 2 copies of *rml* expression cassettes | This study |
| FY3-*rml*-rc*rml* | FY2-*rml* derivative, carrying a *rml* and a rc*rml* expression cassettes | This study |
| FY4-3*rml* | FY3-2*rml* derivative, carrying 3 copies of *rml* expression cassettes | This study |
| FY4-2*rml*-rc*rml* | FY3-2*rml* derivative, carrying 2 copies *rml* a rc*rml* expression cassettes | This study |
| FY5-4*rml* | FY4-3*rml* derivative, carrying 4 copies of *rml* expression cassettes | This study |
| FY5-3*rml*-rc*rml* | FY4-3*rml* derivative, carrying 3 copies of *rml* and a rc*rml* expression cassettes | This study |
| FY6-5*rml* | FY5-4*rml* derivative, carrying 5 copies of *rml* expression cassettes | This study |
| FY6-4*rml*-rc*rml* | FY5-4*rml* derivative, carrying 4 copies of *rml* and a rc*rml* expression cassettes | This study |
| FY7-6*rml* | FY6-5*rml* derivative, carrying 6 copies *rml* expression cassettes | This study |
| FY7-5*rml*-rc*rml* | FY6-5*rml* derivative, carrying 5 copies of *rml* and a rc*rml* expression cassettes | This study |
| Po1f/hp12d-n*rml* | Po1f derivative, carrying n copies of *rml* expression cassettes (n=1-6) | This study |

**Supplemental Table S2. Primers and *lox* site used in this study.**

| **Primer/*lox* site** | **Sequence (5’-3’)** | **Annotation** |
| --- | --- | --- |
| **Primer** |  |  |
| Pox2-F1 | CGCATATGATGCCATCCCACAAGACGAA | PCR for pPOX2 |
| Pox2-R1 | GGAATTGTTATCCGCTCACAATTCCGGATCCGGCGTCGTTGCTTGTGTGATTT | PCR for pPOX2 |
| Cre-F1 | GGAATTGTGAGCGGATAACAATTCCATGTCCAATTTACTGACCGTACA | PCR for *cre* gene |
| Cre-R1 | AGTTGTAAAGAGTGATAAATAGCCCTAGGCTAATCGCCATCTTCCAGCAGGC | PCR for *cre* gene |
| lip2t-F1 | CTGCTGGAAGATGGCGATTAGCCTAGGGCTATTTATCACTCTTTACAACT | PCR for lip2t-*lox*71, lip2t-*lox*66, lip2t-rc*lox*71 |
| lip2t-R1 | GCTAGCACGCGTATAACTTCGTATAATGTATGCTATACGAAGTTATCTCCACCTGTGTCAATCTTC | PCR for lip2t-*lox*71 |
| lip2t-R2 | GGAGTCGCATAAGGGAGAGCTCTAGAGTCGACACGCGTTACCGTTCGTATAATGTATGCTATACGAAGTTATCTCCACCTGTGTCAATCTTC | PCR for lip2t-*lox*66 |
| lip2t-R3 | GGAGTCGCATAAGGGAGAGCTCTAGAGTCGACACGCGTTACCGTTCGTATAGCATACATTATACGAAGTTATCTCCACCTGTGTCAATCTTC | PCR for lip2t-rc*lox*71 |
| Upleu-F1 | AGCATACATTATACGAAGTTATACGCGTGCTAGCCTACGATAAGCAGTCCAATATTCTG | PCR for Upleu-*lox*71 |
| Upleu-R1 | GGCAGGGCCCATAACTTCGTATAATGTATGCTATACGAACGGTAGACAGCAACTACTCCTTTCAC | PCR for Upleu-*lox*71 |
| Upleu-F2 | CAAGCCCGGTCTTACGGCCATC | upstream sequence of Upleu fragment |
| Upleu-F3 | GGTGATGGGAAGAGTCCACTCA | 787-810 bp of Upleu fragment |
| Upleu-R3 | CATACTACAATCACGAGCGCTTC | reverse complementary 820-841 bp of Upleu fragment |
| leu-F2 | CTCATGTTTGACAGCTTATCGCTAGCTACCGTTCGTATAATGTATGCTATACGAAGTTATGATAAGCTGTCAAACATGA | PCR for rc*lox*66-partial *leu2* |
| leu-R2 | TCATCATTTCATTAGCAGGGCAGGGCCCTTTTTATAGAGTCTTATACAC | PCR for rc*lox*66-partial *leu2*, *lox*71-partial *leu2* |
| leu-F3 | CTCATGTTTGACAGCTTATCGCTAGCTACCGTTCGTATAGCATACATTATACGAAGTTATGATAAGCTGTCAAACATGA | PCR for *lox*71-partial *leu2* |
| Ura-F1 | CTGTGCGGTATTTCACACCGCATATGTCAATCCAATTACCCCCCACAAC | PCR for Ura3 |
| Ura-R1 | TTCGTCTTGTGGGATGGCATCATATGGACAAAGGCCTGTTTCTCGG | PCR for Ura3 |
| Apa-R3 | GTACAGGTTCAGGTCCTTTCGCAGC | reverse complementary 616-641 bp of *leu2-270* and *leu2* segments |
| Apa-R6 | AGCGCTATCGAACGTACCCAG | reverse complementary 93-114 bp of *leu2-270* and *leu2* segments |
| KNC-F2 | CATCAACACCGGCCTGTGCACCTAA | 3’ end sequences of *rml* gene |
| SXB-R2 | GCGGATCCACCATTCCTTGCGGCGGCGGTGC | reverse complementary upstream sequence of hp12d promoter |
| qact1 | CTGGCCGAGATCTTACCGAC | qPCR for *act1* gene |
| qact2 | CATCGGGAAGCTCGTAGGAC | qPCR for *act1* gene |
| qRML-F | AAGACCTGGTCCACCCTGATC | qPCR for *rml* gene |
| qRML-F | CGGAGACAGGAGGGTAGGAA | qPCR for *rml* gene |
| ***lox* site** |  |  |
| *lox*P | ATAACTTCGTATAGCATACATTATACGAAGTTAT | wild type |
| *lox*71 | taccgTTCGTATAGCATACATTATACGAAGTTAT | LE mutant |
| *lox*66 | ATAACTTCGTATAGCATACATTATACGAAcggta | RE mutant |
| *lox*72 | taccgTTCGTATAGCATACATTATACGAAcggta | LE+RE mutant |

**Supplemental Table S3.** **Comparison of markerless gene integration tools based on Cre/*lox* and CRISPR-Cas9 systems in *Y. lipolytica*.**

| **Basis** | **Integration manner** | **Marker demand** | **Plasmid recovery** | **Efficiency** | **Reference** |
| --- | --- | --- | --- | --- | --- |
| CRISPR/Cas9 | targeted and makerless integration via DSB and HR | 2 markers | required | 52%-69% on 5 loci, less than 6% on 12 loci | (Schwartz et al., 2017b) |
|  |  |  |  | less than 38% | (Gao et al., 2018) |
|  |  |  |  | more than 80% on 5 loci, less than 30% on 6 loci^†^ | (Holkenbrink et al., 2018) |
| Cre/*lox* | targeted and markerless gene insertion or replacement via HR^*^ | 2 markers | required | 30%-100% on 11 loci^†^ | (Holkenbrink et al., 2018) |
| Cre/*lox* | random and markerless integration | 2 markers | required | 79.2% | (Lv et al., 2019) |
| Cre/*lox* | targeted, markerless and repeated gene insertion via HR | 1 markers | not required | more than 90% | This study |

^*^ The detailed integration manner was not shown in the corresponding reference.

^†^ Target genes were respectively integrated into the same 11 loci via Cre/*lox* and CRISPR-Cas9 tools.

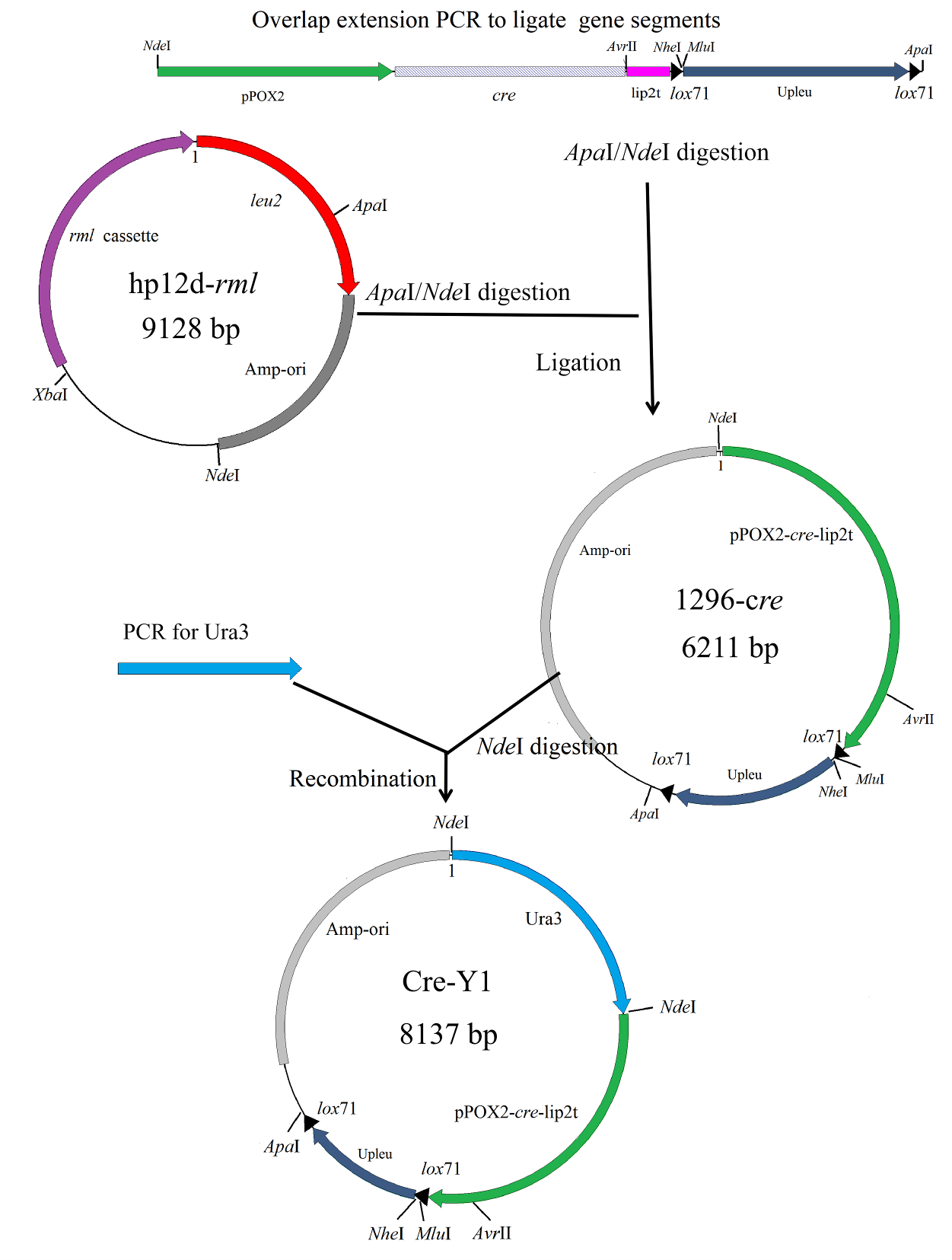

**Supplemental Fig. S1.**  Process diagram for constructing plasmid Cre-Y1.

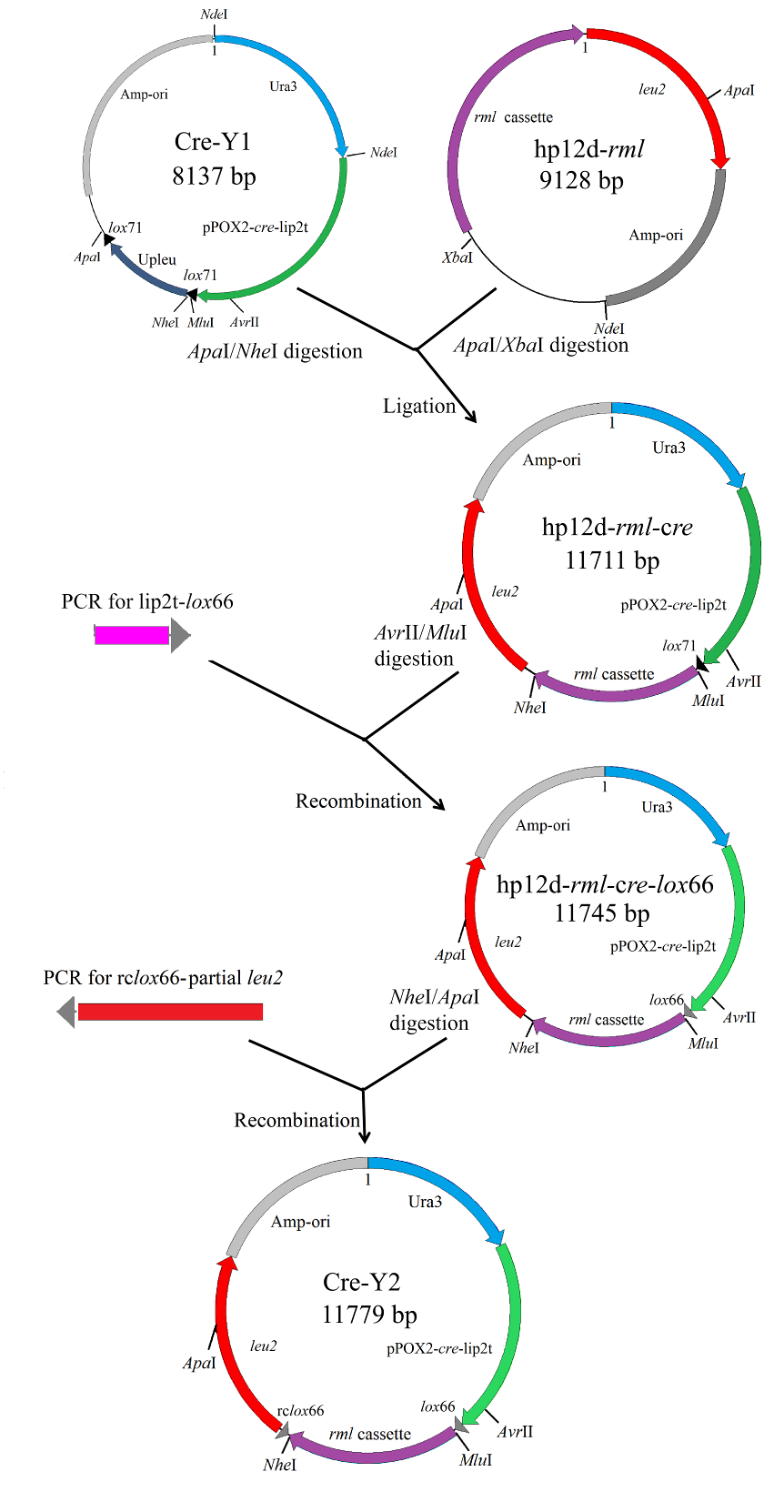

**Supplemental Fig. S2.** Process diagram for constructing plasmid Cre-Y2.

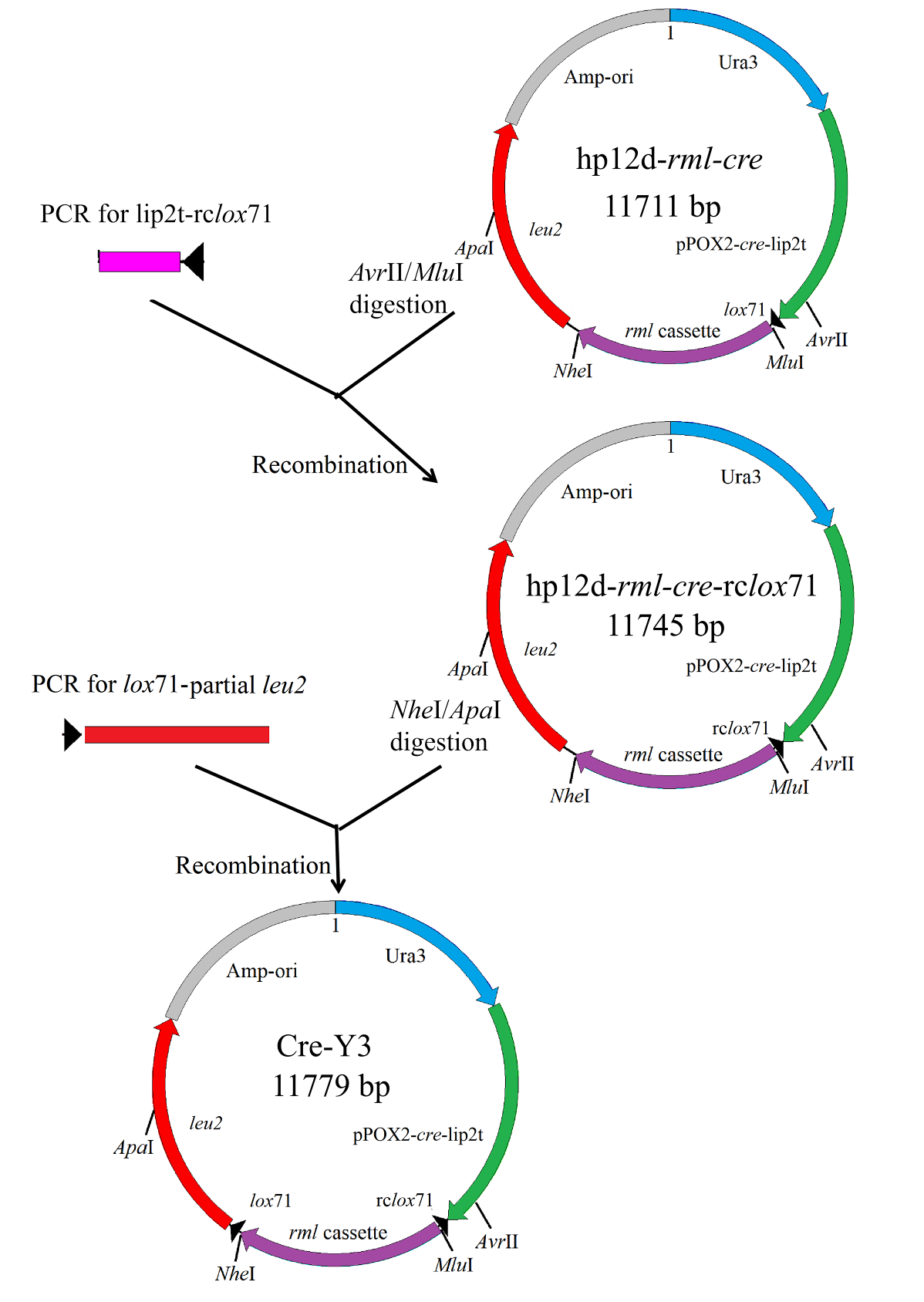

**Supplemental Fig. S3.**  Process diagram for constructing plasmid Cre-Y2.

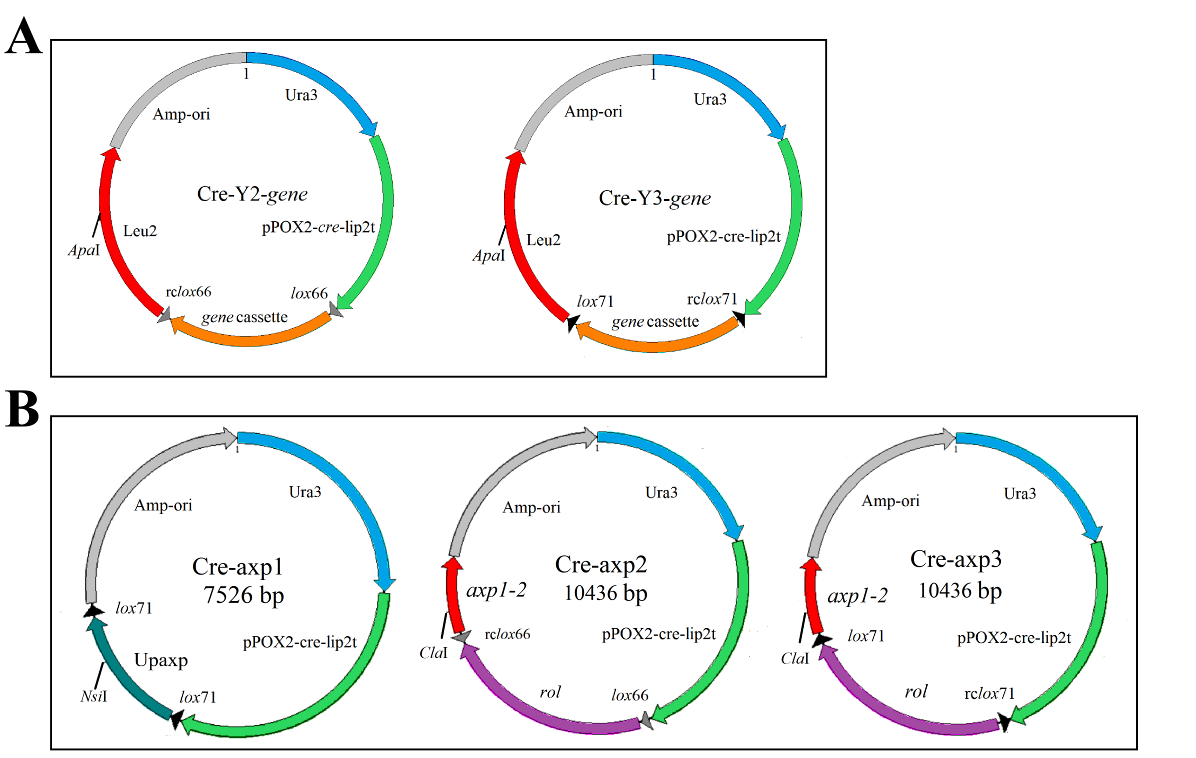

**Supplemental Fig. S4.**  (A) Plasmids for various gene integrations in *leu2* locus of Po1f. These plasmids were derived from Cre-Y2 or Cre-Y3 and carried one of the following genes: *ire1*, *kar2*, *pdi*, *sls1*, *hac1,* and *vgb*. All plasmids were linearized by *Apa*I at homologous fragment *leu2*. (B) Plasmids for *rol* gene integration in *axp1-2* locus of Po1h. The plasmids Cre-axp1, Cre-axp2, and Cre-axp3 were respectively derived from Cre-Y1, Cre-Y2, and Cre-Y3. The Upaxp is the upstream segment of *axp1-2* gene (about 1000 bp) and *axp1-2* is designed as the homologous fragment of *axp1-2* in Po1h, and their functions are the same as Upleu and *leu2* in the original genetic tool.

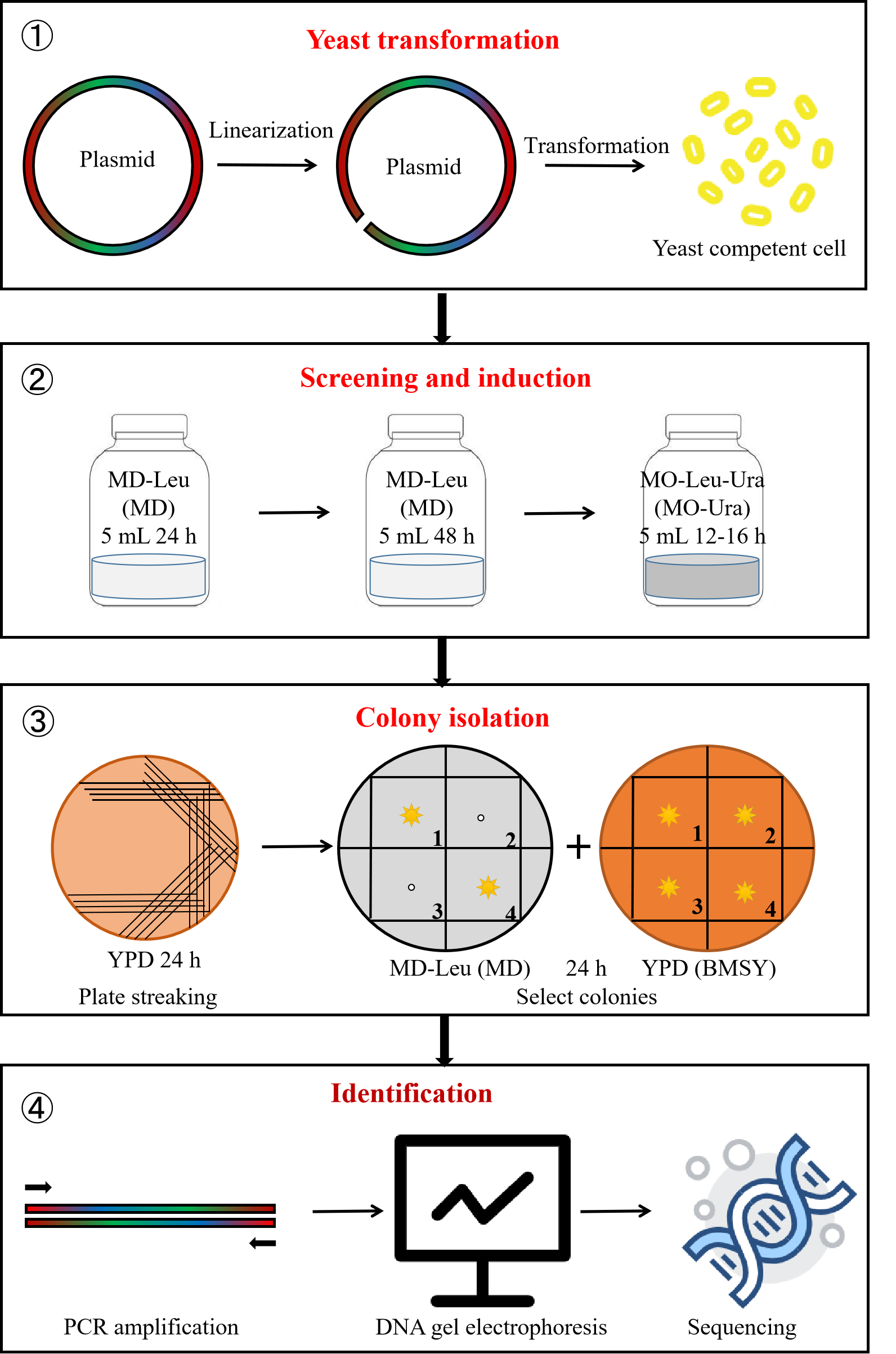

**Supplemental Fig. S5.** Illustration of the gene integration process employed in this study. The integration process included yeast transformation, screening and induction, colony isolation, and identification. First, the plasmids were linearized and used to transform yeast competent cells. Then, the transformants were screened in 5 mL of MD-Leu (MD) medium for 24 h, and rescreened in 5 mL of MD-Leu (MD) liquid medium for 48 h. Subsequently, 100 μL of the cell suspension solutions were induced in 5 mL of MO-Leu-Ura (MO-Ura) liquid medium. Afterwards, single colony was obtained by plate streaking on YPD medium and phenotype identification on YPD (BMSY) and MD-Leu (MD) media. The selected positive colonies were further confirmed via PCR amplification and DNA sequencing.

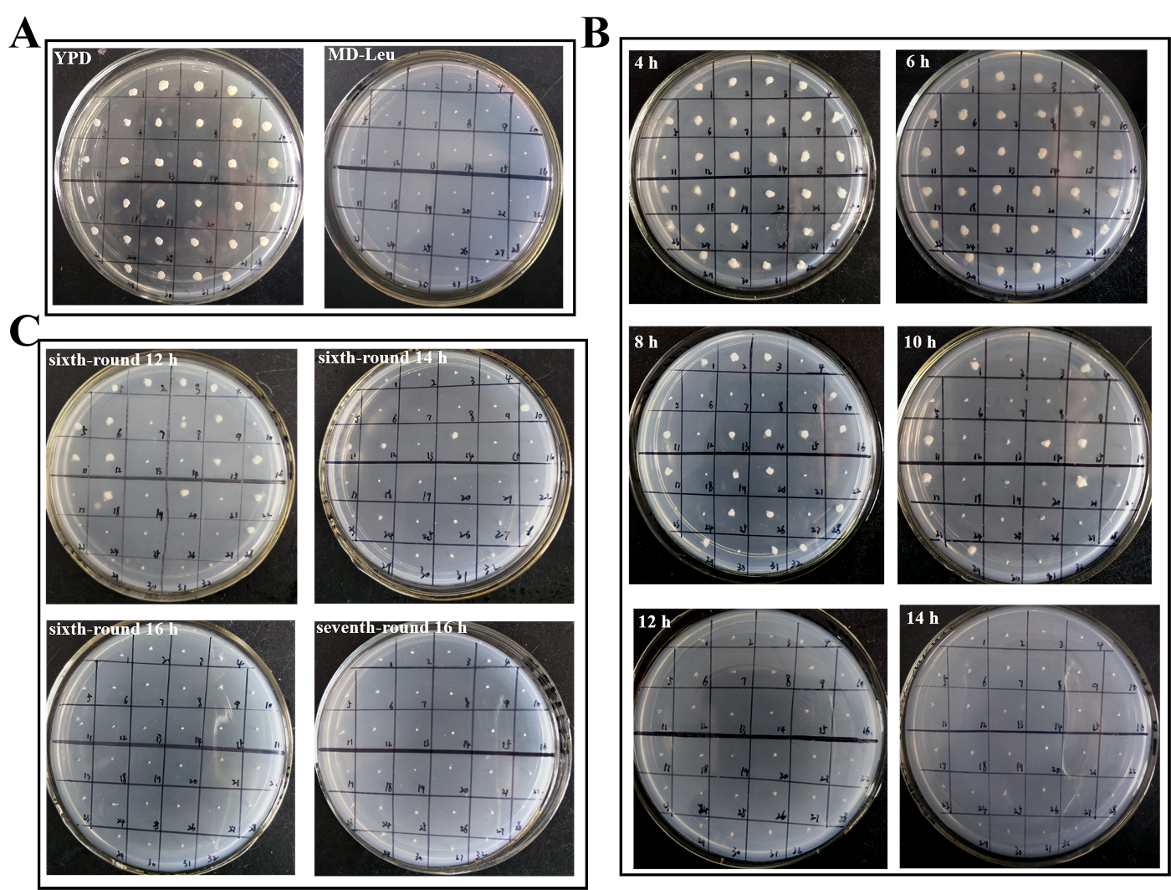

**Supplemental Fig. S6.** (A) Isolation of positive colonies for FY1. A total of 32 recombinants were randomly selected and inoculated onto YPD and MD-Leu media, and none of them could grow on MD-Leu medium. (B) Positive colonies induced for 4–14 h in the second-round genome integration. The corresponding BMSY plates are not shown. (C) Positive colonies induced for 12–16 h in the sixth and seventh rounds of integration. The corresponding BMSY plates are not shown.

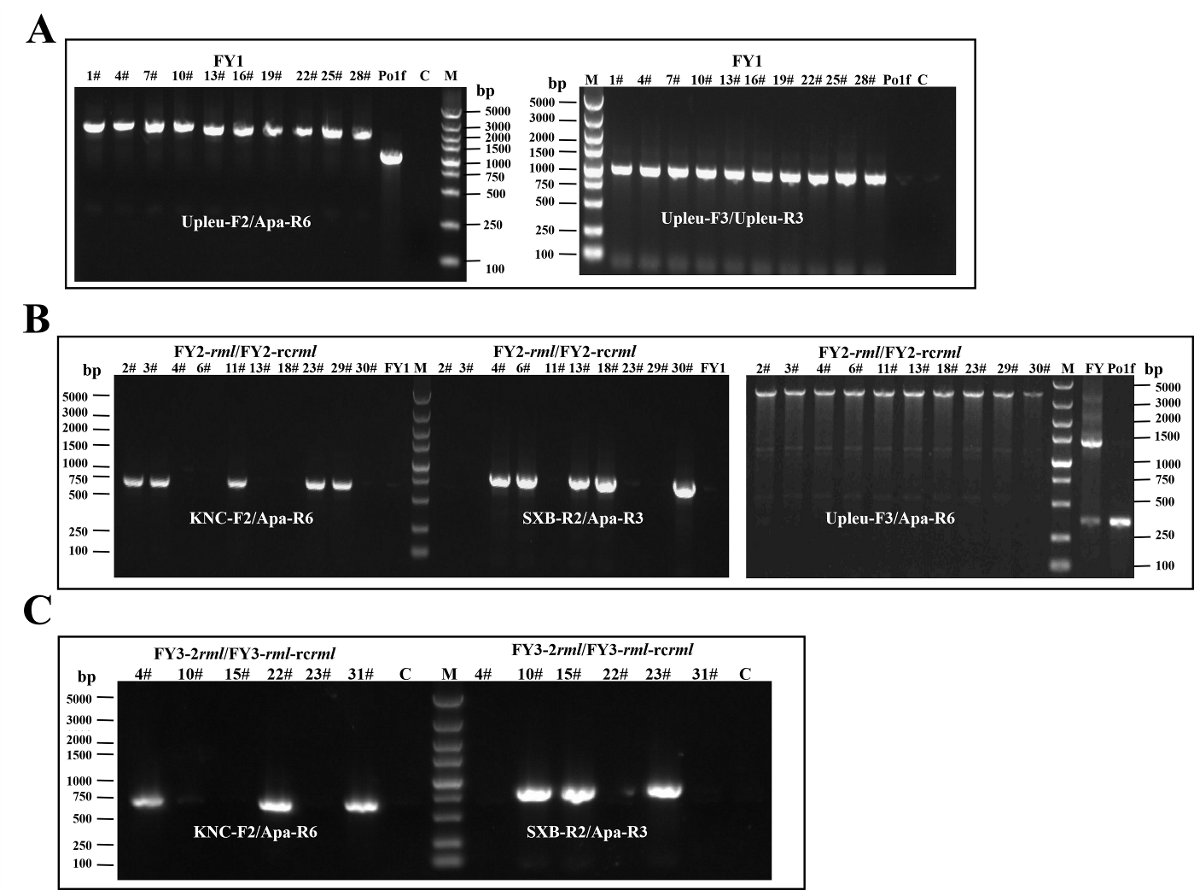

**Supplemental Fig. S7.** PCR amplification to identify positive colonies in the first–third rounds of integration. Lane M, Marker; lane C, ddH_2_O. (A) PCR product of about 2400 bp (corresponding to Upleu-*lox*71-Upleu-partial *leu2-270* in FY1 genome) and about 1300 bp (corresponding to Upleu-partial *leu2-270* in Po1f genome) were respectively amplified by primer pair Upleu-F2/Apa-R6. The PCR product corresponding to Upleu-*lox*71-Upleu (about 900 bp) was obtained using primers Upleu-F3/Upleu-R3 in FY1; however, no product was obtained in Po1f. (B) The fragment partial *rml*-rc*lox*66-partial *leu2* (about 650 bp) was amplified with the primer pair KNC-F2/Apa-R6 only in FY2-*rml*. The segment partial rc*rml*-rc*lox*66-partial *leu2* (about 800 bp) was cloned by SXB-R2/Apa-R3 only in FY2-rc*rml*. Different PCR products were obtained in four types of strains via amplification with Upleu-F3/Apa-R6: Upleu-*lox*72-*rml/*rc*rml*-rc*lox*66-partial *leu2* (about 3500 bp) in FY2-*rml*/FY2-rc*rml*, Upleu-*lox*71-Upleu-partial *leu2* (about 1300 bp) in FY1, and Upleu-partial *leu2* (about 300 bp) in FY1 and Po1f. (C) PCR products of about 650 bp (corresponding to partial *rml*-rc*lox*66/*lox*71-partial *leu2*) were amplified with primer pair KNC-F2/Apa-R6 only in FY3-2*rml*, and DNA fragments of about 800 bp (corresponding to partial rc*rml*-rc*lox*66/*lox*71-partial *leu2*) were cloned using SXB-R2/Apa-R3 in FY3-*rml*-rc*rml*.

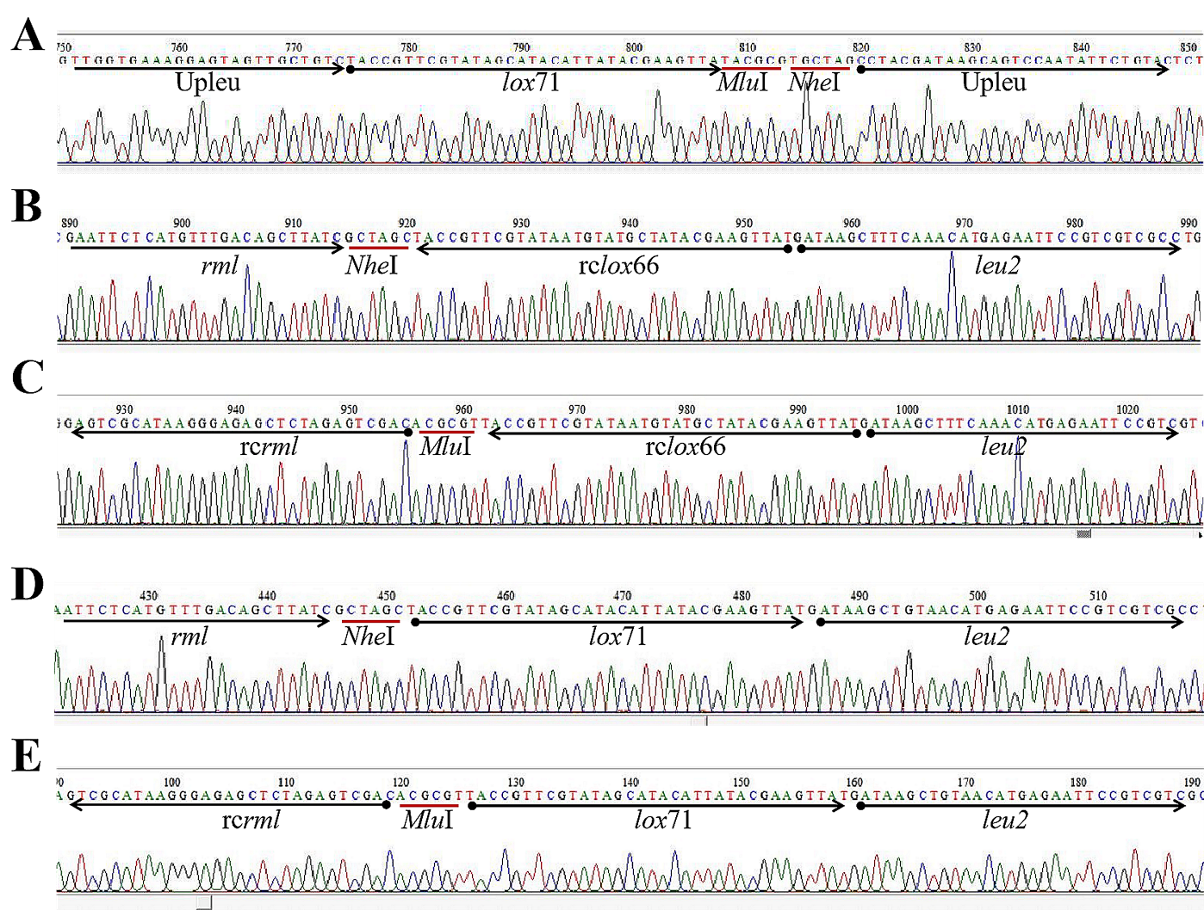

**Supplemental Fig. S8**. Partial sequencing result of PCR product in this study. (A) Upleu-*lox*71-Upleu from FY1. (B) *rml*-rc*lox*66-*leu2* from FY2-*rml*. (C) rc*rml*-rc*lox*66-*leu2* from FY2-rc*rml*. (D) *rml*-*lox*71-*leu2* from FY3-2*rml*. (E) rc*rml*-*lox*71-*leu2* from FY3-*rml*-rc*rml*.

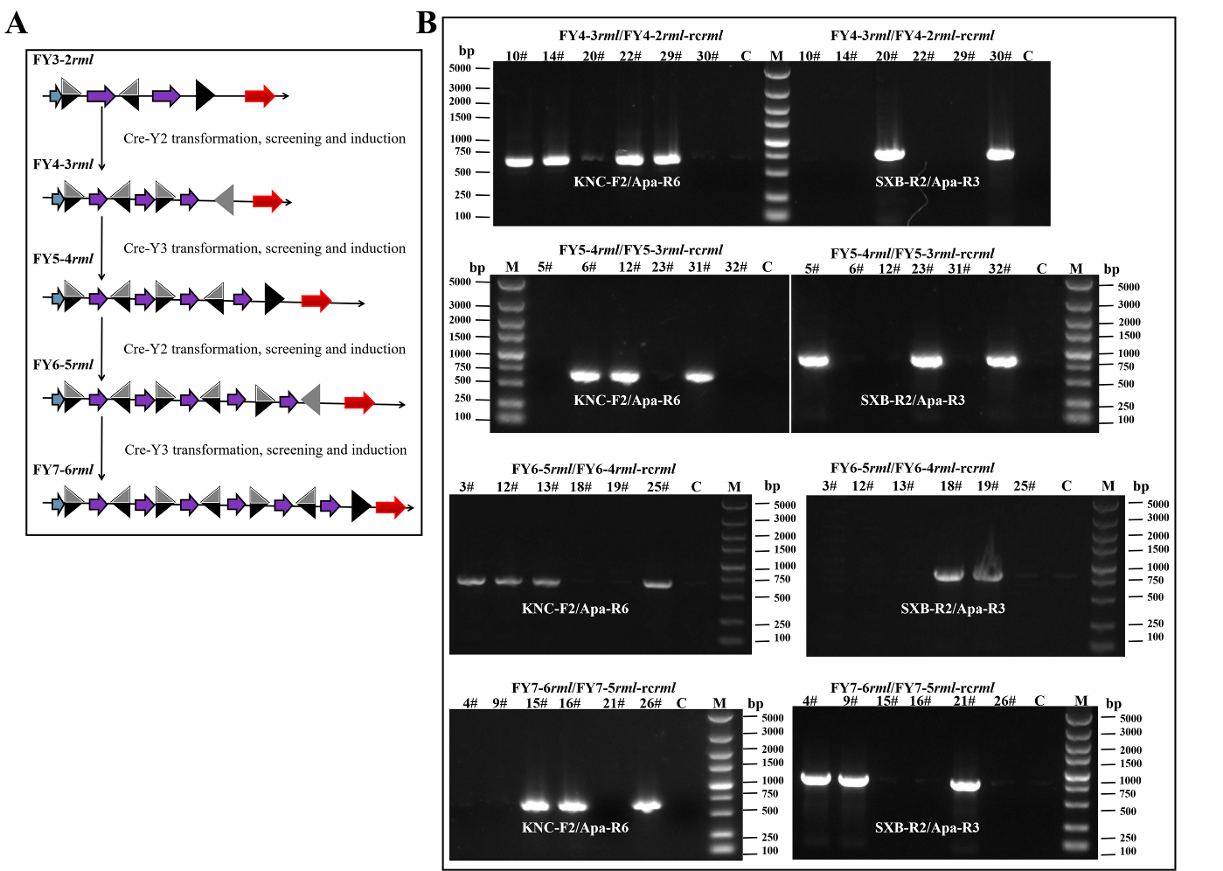

**Supplemental Fig. S9.** (A) Schematic diagram of the fourth–seventh rounds of integration. The recombinant strains harboring rc*rml* expression cassettes were not present. (B) PCR amplification to identify positive colonies in the fourth–seventh rounds of integration. The PCR products on the left (about 650 bp, corresponding to partial *rml*-rc*lox*66/*lox*71-partial *leu2*) were amplified with primer pair KNC-F2/Apa-R6, and the DNA fragments on the right (about 800 bp, corresponding to partial rc*rml*-rc*lox*66/*lox*71-partial *leu2*) were cloned using SXB-R2/Apa-R3. Lane M, Marker; lane C, ddH_2_O.

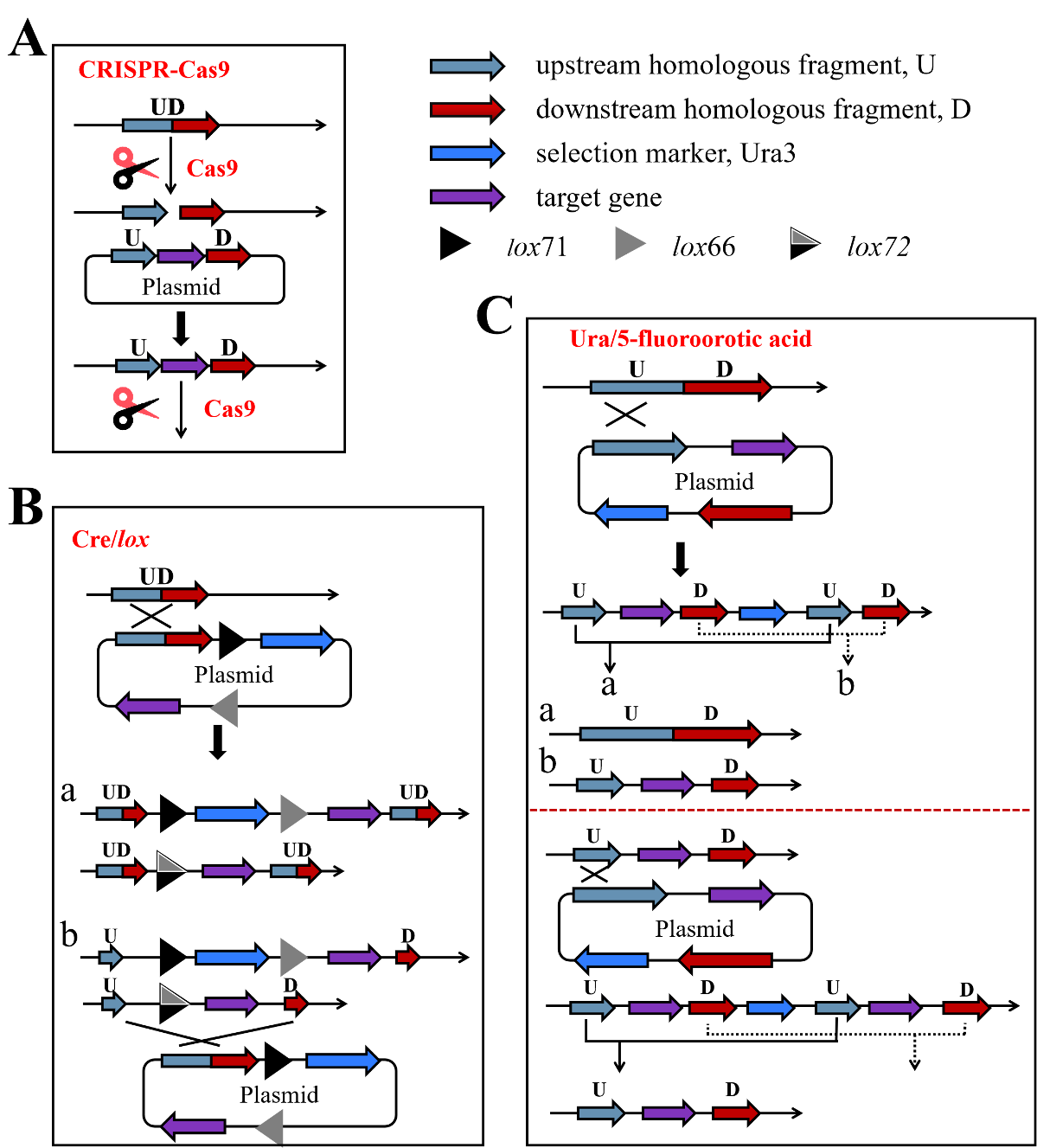

**Supplemental Fig. S10.** Repeated, targeted and markerless gene integration via the existing genetic tools based on CRISPR-Cas9 system, Cre/*lox* system, and Ura/5-fluoroorotic acid resistance method. The integrated states are similar if the plasmids in this figure are replaced by DNA fragments carrying target genes. (A) Cas9 protein and sgRNA can cause DSB in the homologous fragment UD, and the target gene can be integrated into UD locus via HR repair. However, the target gene may be broken if the same sgRNA is used for subsequent gene integration. Thus, more integration rounds require additional plasmids harboring new sgRNAs. (B) Recombinant plasmid carrying target gene is integrated into homologous fragment UD via gene insertion (a) or gene replacement (b), and the Ura marker can be removed by deletion reaction between *lox* sites. Gene insertion event is highly frequent and can retain two UD fragments, resulting in gene redundancy in the genome. On the contrary, gene replacement event is more suitable, but infrequent. Nonetheless, target gene will be replaced if the UD fragment is used for second-round integration. Therefore, further integration requires different homologous fragments. (C) Plasmid harboring target gene is inserted into the homologous fragment U. Although two kinds of recombination reactions (a and b) may occur via Ura/5-fluoroorotic acid resistance, only reaction b produces target strain. Moreover, the recombination reactions will generate reversible strains when the same plasmid is integrated in the above-mentioned target strain (the HR of target gene segments is not present). Hence, new homologous fragments should be employed for additional gene integration.
